# Tracking acquisition of cognitive maps in early- and late-onset blindness

**DOI:** 10.64898/2026.05.18.726055

**Authors:** Maxime Bleau, Quentin Dessain, Laurence Dricot, Joseph Paul Nemargut, Ron Kupers, Maurice Ptito

**Author notes:** **Corresponding authors**: Maxime Bleau, Maurice Ptito, **Email:**.

## Abstract

Cognitive maps encode spatial relationships between locations and support flexible navigation. However, how these mental representations form in blindness remains unclear. Here, we introduce a multisensory virtual navigation paradigm that monitors non-visual cognitive map formation over time. Sixteen early blind (EB), 17 late blind (LB), and 29 sighted controls (SC) learned the layout of a tactile maze and, in a virtual version, repeatedly performed pointing (judgment of relative directions) and navigation (reaching locations) tasks. Thus, this experiment tracked both functional and spatio-cognitive components of cognitive map formation across multiple learning stages and showed that EB accumulated knowledge more slowly than SC. In addition, both EB and LB had reduced pointing accuracy under viewpoint changes, indicating difficulty translating cognitive maps into first-person perspectives. Yet, navigation and pointing improved in all groups, and a subset of EB and LB achieved expert-level performance. In sum, this study provides a scalable framework for tracking the temporal dimension of cognitive map formation in blindness and other neurological conditions, informing how spatial knowledge is acquired and translated into navigation behavior. Importantly, it suggests that allocentric cognitive map formation in blindness is an experience-dependent process that varies across time and individuals. Observed spatial disadvantages in both blind groups may result from adaptive strategies that constrain the use of allocentric cognitive maps, independent of prior visual experience.

## Introduction

Cognitive maps are internal representations of the environment that guide goal-directed behaviors, including spatial navigation (1, 2). These representations are typically described as “allocentric” (e.g., world-centered, independent of the individual’s viewpoint) rather than “egocentric” (e.g., centered on the individual’s viewpoint) (1, 3). However, cognitive map formation remains difficult to characterize and monitor over time, particularly in populations for whom standard experimental paradigms are poorly suited. This is especially true for individuals living with severe sensory limitations, such as blindness. Because vision is the primary sensory channel supporting spatial cognition in humans (4), individuals with blindness face significant challenges in building cognitive maps, which affect their orientation abilities, independence, and quality of life (5, 6).

To date, no clear consensus exists on the extent to which blindness affects cognitive mapping abilities (7, 8). Numerous studies report that early blind (EB) individuals have normal or supranormal abilities in real-world navigation tasks. These tasks include learning new routes, retracing and combining routes, taking shortcuts, pointing to locations, reproducing maze layouts, and estimating distances between locations after exploration (9–16). Other studies, however, report disadvantages for EB when tasks require allocentric processing, including cognitive map formation. Several studies show that EB are less proficient in acquiring and using survey-based spatial knowledge of large-scale environments (17–20). This translates into less accurate estimation of directions and distances (4, 21–27) and compressed internal spatial representations (28). These difficulties become particularly apparent when tasks require mental rotation or inference of spatial relationships (17, 29, 30).

Nonetheless, individuals with blindness can form cognitive maps of complex environments (7, 31–34) and become expert navigators with O&M training and lifelong, repeated practice (35–37). There is thus no doubt that the functional impacts of blindness are multifactorial, likely stemming from complex interactions between personal and environmental factors (8, 38, 39). Investigating these interindividual differences in cognitive map-based spatial navigation is as important for understanding and training spatial cognition in blind individuals as in sighted individuals (40–44). However, assessing the cognitive map’s accuracy and flexibility (the spatio-cognitive component) alongside the navigation behavior it supports (the functional component) in blindness remains challenging, particularly while the cognitive map is still forming.

Across the literature, non-visual cognitive maps are assessed in variable ways. Discrepancies have been attributed in part to the used environments and paradigms (7, 45): reports of preserved spatial abilities come from simple to moderately complex layouts, while reports of impairments come from complex routes and layouts (7, 8, 45), or from paradigms that rely on strategies typical of visually guided navigation (e.g., triangle completion) but suboptimal for individuals relying on non-visual sensory input (35, 46). Furthermore, measures probing spatial knowledge, including Euclidean pointing, judgments of relative direction (JRD), map reconstruction, and shortcutting (31, 47–50), are often administered once, after learning (7, 16, 49, 51). However, the temporal dimension of cognitive map formation is important: individuals often differ more in the rate at which spatial knowledge accumulates or decays (40–44, 52, 53), and these phenomena follow non-linear curves that are estimable and comparable across groups (53, 54), as used to characterize spatial memory change in aging and neurodegeneration (52, 53, 55–58).

This learning rate is especially informative in blindness, where survey knowledge improves with training and access to spatial information (7, 21, 31, 36, 59), but group contrasts are commonly reported without establishing how each group learned (7, 8). Tracking cognitive map formation over time within participants, using a standardized measure, could address this gap and help answer long-standing questions about knowledge acquisition without vision (8, 35, 46, 60). However, robust paradigms that track this temporal aspect of cognitive map formation in blindness, in a way that is sensitive to interindividual variability, are lacking. Consequently, the formation of non-visual cognitive maps and their use during navigation remain poorly understood, despite their high importance for individuals’ autonomy (8, 46).

In the present study, we introduce a novel “spatial learning” paradigm to track the formation of non-visual cognitive maps over time in early blind (EB), late blind (LB), and blindfolded sighted individuals. This protocol combines 3D-printed tactile maps (61, 62) with virtual environments, both of which enable blind individuals to build cognitive maps (32, 63–67) that can translate into real-world scenarios (11, 32, 63, 64, 66–71) and even surpass those built through direct exploration (31, 59, 72–74). Using these tools, we quantified cognitive maps over time based on navigation performance and pointing accuracy (via JRD). Whereas the former reflects how cognitive maps support navigation behavior (functional component), the latter reflects how they are flexibly used under different viewpoint changes (spatio-cognitive component). This study therefore establishes the time course of cognitive map formation across learning and disentangles how visual experience and adaptation to permanent vision loss contribute to this process.

## Results

Sixty-two participants took part in the study, including 16 EB (9M, 7F, mean age = 40.8 ± 13.2 years), 17 LB (10M, 7F, mean age = 45.0 ± 11.0 years), and 29 blindfolded sighted controls (SC; 15M, 14F, mean age = 35.4 ± 12.5 years). The experiment consisted of twelve blocks of the “spatial learning” paradigm (see Methods), each consisting of three subtasks: 1) a tactile exploration phase (60s, once per block) to form and refine their cognitive map of the maze; 2) a pointing phase (12s, 8 times per block) in which participants were given a starting point and destination and then instructed to judge the relative direction (JRD) of the destination (from the viewpoint of the starting position); and 3) a virtual navigation phase (maximum of 25s, 8 times per block) in which participants virtually moved through the maze to reach the destination. Hallways in the virtual maze followed a “tile” system: participants could move from tile to tile, and distance was measured in tiles (Fig. 1a).

**Figure 1.**
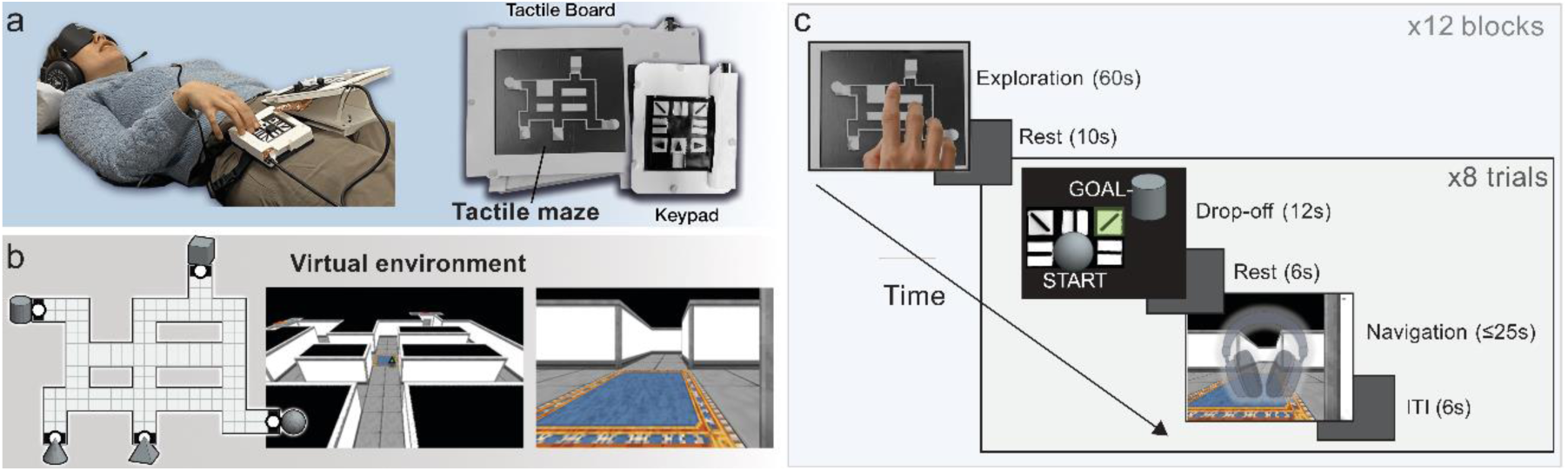
Apparatus and experimental procedures. **a.** The apparatus consisted of 1) a tactile board on which the tactile maze was placed, and 2) a custom keyboard with arrow keys (for navigation) and five oriented bars (for pointing). **b.** The virtual environment recreates the tactile maze. The figure shows the underlying layout and tile system, along with the top and first-person views of the maze in engine. **c.** The protocol consisted of 12 repeated blocks, each comprising one exploration phase and 8 trials (pointing + navigation). Abbreviations: ITI, Intertrial Interval.

To ensure reliance on internal spatial representations, no salient landmarks were provided during virtual navigation, making success dependent on participants’ memory (their cognitive map) and their ability to improvise routes between destinations. Learning curves to monitor cognitive map formation throughout the experiment were defined according to the average performance for each block, resulting in learning curves of 12 mean data points for each group. Learning curves were calculated for three main performance measures: average navigation performance (whether the destination was reached, in %), navigation pace (in seconds per tile; in successful trials only), and average pointing accuracy in the JRD task (pointing answer vs. real direction; for more detailed information, see Methods). Figure 1 provides a detailed representation of the apparatus and experimental procedures.

### Slower spatial knowledge acquisition in EB

Across the entire experiment, SC achieved an average navigation performance of 67.06 ± 19.34%; EB, 55.46 ± 24.04%; and LB, 54.88 ± 23.47% (fig. 2a). During the last three blocks of the experiment, all groups showed improved performance: EB reached 62.68 ± 28.54% (+31.62% from baseline), LB 70.78 ± 23.60% (+44.77% from baseline), and SC 80.80 ± 19.68% (+48.75% from baseline), suggesting that all groups formed a cognitive map of the maze and used it successfully for navigation, with EB improving the least. We fitted linear mixed-effects (LME) models to trial-level data, with participants as a random effect and the learning curve modeled as an exponential curve (see methods SI Appendix). Omnibus tests were then performed to determine if groups differed in terms of i) baseline performance, ii) average performance (level), and iii) improvement rates (group × block interaction; slope of the learning curve). The test showed that groups differed significantly in improvement rates (χ²(2) = 8.77, p = .012; Fig. 2b) but did not differ at baseline (Wald χ²(2) = 0.25, p = .881; Fig. 2b) or in average performance (χ²(2) = 1.18, p = .555; Fig. 2a). According to the LME models, all three groups improved reliably across the experiment (EB β = 34.72, z = 6.59; LB β = 41.31, z = 8.05; SC β = 53.90, z = 13.74; all p < .001; Fig. 2b). Decomposing the interaction, EB improved more slowly than SC (β = 19.18, z = 2.92, p = .004, p_holm_ = .011). In contrast, improvements in LB did not differ from those in EB (β = 6.59, z = 0.90, p = .371, p_holm_ = .371) or in SC (β = 12.59, z = 1.95, p = .051, p_holm_ = .103). For a complete report of the LME analyses, including confidence intervals and effects of age and sex covariates, see SI Appendix. Taken together, these results suggest slower spatial knowledge accumulation in EB and/or difficulties in translating allocentric spatial representations into efficient navigation behavior, while LB exhibit abilities intermediate between EB and SC.

**Figure 2.**
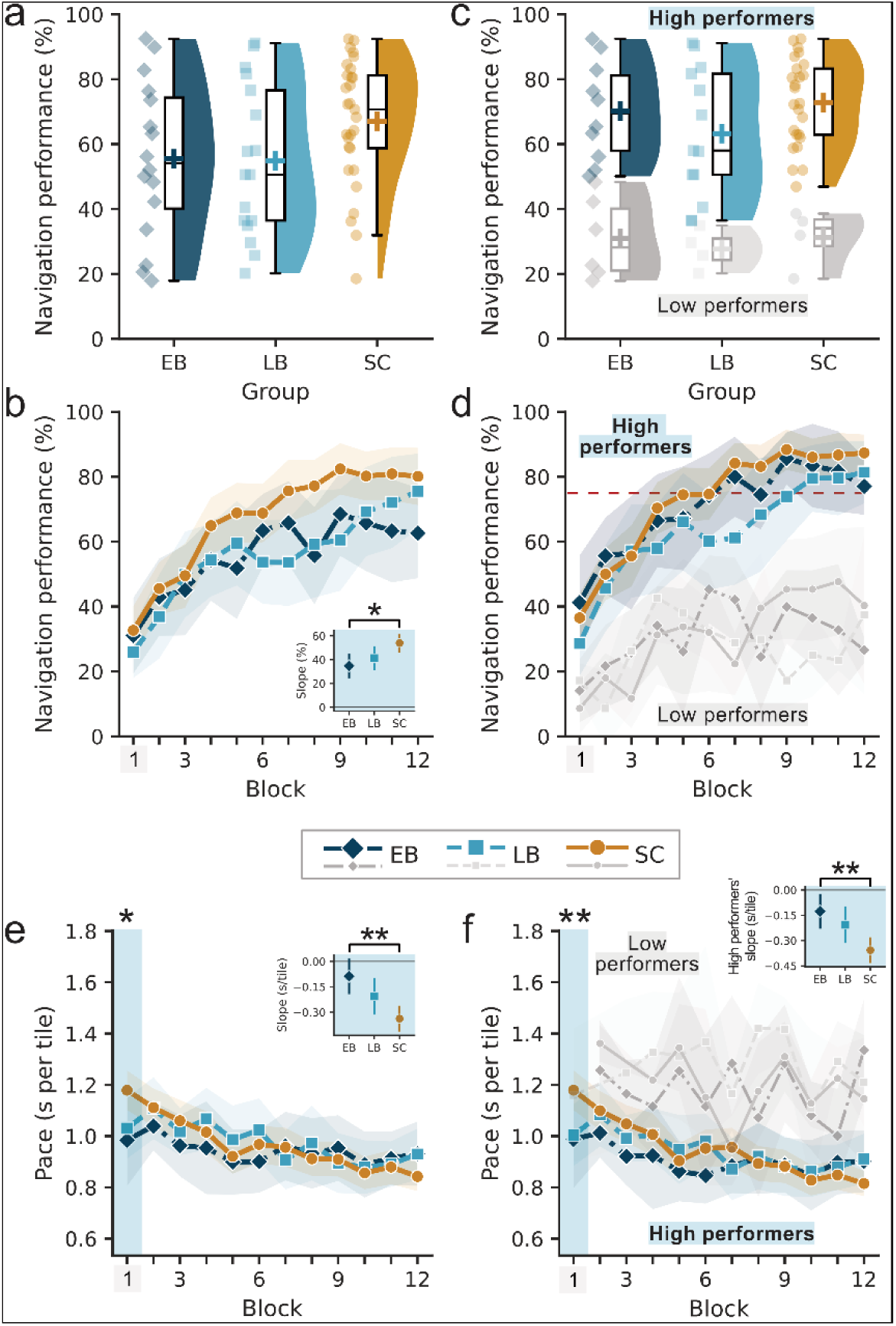
Average navigation performance and learning curves. **a.** Average navigation performance for each participant (individual data points) and the overall distribution; group averages are shown with crosses. **b.** Improvements in navigation performance across the 12 blocks and the significant difference in slope between EB and SC. **c.** Average navigation performance for high (bold, colored) and low performers (thin, greyscale). **d.** Improvements in navigation performance across the 12 blocks for high (bold, colored) and low performers (thin, greyscale), against the 75% learning threshold (red dashed line). **e.** Improvements in pace across the 12 blocks, showing a significant difference in slope between EB and SC and a significant difference at baseline (blue rectangle), where EB and LB were generally faster than SC. **f.** Improvements in pace across the 12 blocks for high (bold, colored) and low performers (thin, greyscale), and the difference in slope between EB and SC high performers. For each error bar plot displaying learning rates, areas around group lines depict confidence intervals (95%). †, p<.10; *, p<.05; **, p<.01; ***, p<.001. Abbreviations: EB, Early Blind; LB, Late Blind; s, seconds; SC, Sighted Controls.

### High and low performers

We separated participants into high and low performers. Low performers were defined as those who did not reach a 75% block performance at least once, while high performers (the rest of the sample) eventually passed this threshold (see Fig. 2d). Consequently, low-performing participants were those who could not learn the maze well enough to navigate between locations successfully within the time limits. According to this criterion, low performers included 6/16 EB (37.50%), 4/17 LB (23.53%), and 4/29 SC (13.79%). Because the smallest expected cell was 3.61, we performed a permutation test on the chi-square statistic and found no significant differences among these proportions (χ²(2) = 3.33, p = .192, 20,000 permutations, Cramér’s V = 0.232). Nonetheless, the EB group showed a qualitatively higher proportion of low performers. As a supplementary classification, we also defined participants who reached the 75% learning threshold early (e.g., in blocks 1 or 2) and maintained an average performance of ≥ 90% as expert map-users. Expert map-users were present across all groups: 2/16 EB (12.50%), 2/17 LB (11.76%), and 3/29 SC (10.34%), suggesting that highly successful allocentric, cognitive map-based navigation is attainable with or without visual experience. Because of the limited number of expert map-users, we did not include this subdivision in the analysis, which was limited to the high- and low-performer classification.

We then conducted the same LME analyses separately for low and high performers. Among high performers, the exponential block term provided the best fit (see SI Appendix), and the group × block interaction was not significant (χ²(2) = 3.28, p = .194). The groups differed neither in terms of overall performance (χ²(2) = 1.44, p = .486) or baseline performance (χ²(2) = 1.54, p = .464), but all three groups improved (EB β = 43.09, LB β = 48.59, SC β = 56.27, all p_holm_ < .001). For low performers, the best fit was a quadratic block term (see SI Appendix). The omnibus tests revealed a group × block interaction (χ²(4) = 11.06, p = .026), but no differences in overall performance (χ²(2) = 0.17, p = .920), nor in baseline performance (χ²(2) = 0.97, p = .615). The LME models revealed that EB and SC improved and, then, decelerated, while LB did not (EB: linear β = 7.00, z = 3.72, p_holm_ = .001, curvature β = −0.531, z = −3.14, p_holm_ = .008; SC: linear β = 5.84, z = 2.61, p_holm_ = .037, curvature β = −0.235, z = −1.20, p_holm_ = .235; LB: linear β = 4.55, z = 1.91, p_holm_ = .169, curvature β = −0.350, z = −1.57, p_holm_ = .235). However, no pairwise contrast was significant (all p_holm_ = 1.00), mainly due to the small sample of low-performing participants. Taken together, these results suggest that the whole-sample group × block interaction may have been driven by a larger proportion of low-performing EB participants. More information is available in the SI Appendix.

### Navigation efficiency (pace towards destination)

We estimated navigation efficiency as pace (seconds per tile; total time / start-goal Euclidean distance; see methods) in successful trials. Because 3,054 of 5,835 trials were successful, we analyzed this measure as a selected subset. First, the equivalence between the three groups was tested: EB failed to reach the destination on 53.6 ± 27.5% of trials, LB on 53.1 ± 25.9%, and SC on 41.5 ± 23.3%; these proportions did not significantly differ (F(2,57) = 1.03, p = .365, η²_p_ = 0.035). Second, we weighted each observed trial by the inverse of its modeled probability of arrival, estimated from all 5,835 trials using group, block, start-goal Euclidean distance, age, and sex (see methods and SI Appendix).

An exponential form best described the learning curve for pace (see SI Appendix). The omnibus tests revealed a significant group × block interaction (χ²(2) = 14.33, p < .001), with a group difference at baseline level (Wald χ²(2) = 7.08, p = .029), but no overall group difference (χ²(2) = 0.76, p = .685). Decomposing the interaction, the model shows that SC improved faster than EB (β = −0.277, z = −3.75, p < .001, pholm < .001), with LB intermediate between SC and EB (pholm = .128 and .128, respectively). At baseline, SC were slower than EB (β = 0.198, z = 2.52, p = .012, pholm = .035), and LB were intermediate between SC and EB (pholm = .207 and .223, respectively). Consistently, LB and SC improved (LB: β = −0.235, z = −4.03, p < .001, pholm < .001; SC: β = −0.371, z = −8.41, p < .001, pholm < .001), whereas EB did not (β = −0.094, z = −1.61, p = .108, pholm = .108). More information is available in the SI Appendix.

The same analyses (with the logarithmic form as the best fit; see SI Appendix) were conducted for high and low performers (Fig. 2f). The omnibus tests revealed similar patterns among high performers (2,811 successful trials): it showed a group × block interaction (χ²(2) = 13.90, p = .001), a difference at baseline (Wald χ²(2) = 9.79, p = .007), but no difference in overall performance (χ²(2) = 1.54, p = .462). SC improved faster than EB (β = −0.095, z = −3.54, p < .001, p_holm_ = .001) and LB (β = −0.061, z = −2.26, p = .024, p_holm_ = .048), but EB and LB did not differ from each other (β = −0.034, z = −1.09, p_holm_ = .276). At baseline, SC were slower than both EB (β = 0.232, z = 2.78, p = .005, p_holm_ = .016) and LB (β = 0.136, z = 2.33, p = .020, p_holm_ = .040), while LB and EB did not differ (β = 0.096, z = 1.16, p = .247, p_holm_ = .247). Contrary to the whole sample, however, high performers in all three groups became faster (EB: β = −0.049, z = −2.26, p = .024, p_holm_ = .024; LB: β = −0.083, z = −3.72, p < .001, p_holm_ < .001; SC: β = −0.144, z = −9.56, p < .001, p_holm_ < .001). Among low performers, only 243 trials were successful, rendering the analysis uninformative. Taken together, these results suggest that the lack of improvement in EB may be driven by participants who never reached the learning criterion. In contrast, high-performing EB participants improved in pace. More information is available in the SI Appendix.

### Altered allocentric coding in early blindness

Across all trials, allocentric cognitive map formation was also monitored by the accuracy with which participants pointed to destinations during the pointing task. In each trial, pointing accuracy was measured as participants’ pointing scores (%), which indicate how close participants’ direction estimates were to the actual direction (the angle between the starting location and the destination). Consequently, accurate cognitive maps would lead to more precise pointing, whereas lower pointing scores indicate distortions in participants’ mental representations. After excluding invalid responses (74 trials; see Methods and SI Appendix), pointing estimates were provided in 5,295 trials: per group, pointing estimates were provided on 91.9% (EB), 90.9% (LB), and 92.7% (SC) of trials.

On average, SC (average pointing scores of 84.88 ± 6.74%) had better allocentric pointing abilities than EB (76.70 ± 7.84%), and LB (80.30 ± 7.09%) were intermediate between SC and EB (Fig. 3a). This was confirmed by the LME analysis (with pointing scores described by a power-law approach to asymptote, see SI Appendix): the omnibus tests revealed a significant difference in overall pointing scores (χ²(2) = 10.57, p = .005), with no group × block interaction (χ²(2) = 3.95, p = .138), and no difference at baseline (Wald χ²(2) = 3.56, p = .169). EB were significantly less accurate than SC (β = 7.14, z = 3.33, p < .001, p_holm_ = .003), while LB were intermediate (vs SC: β = 4.02, z = 1.87, p = .061, p_holm_ = .122; vs EB: β = 3.12, z = 1.31, p = .191, p_holm_ = .191; see Fig. 3a-b). While improvements did not differ, all groups improved in pointing accuracy across the experiment (EB: β = 7.29, z = 2.14, p = .032, p_holm_ = .032; LB: β = 16.23, z = 4.95, p < .001, p_holm_ < .001; SC: β = 14.50, z = 5.83, p < .001, p_holm_ < .001; see Fig. 3b). More information is available in the SI Appendix.

**Figure 3.**
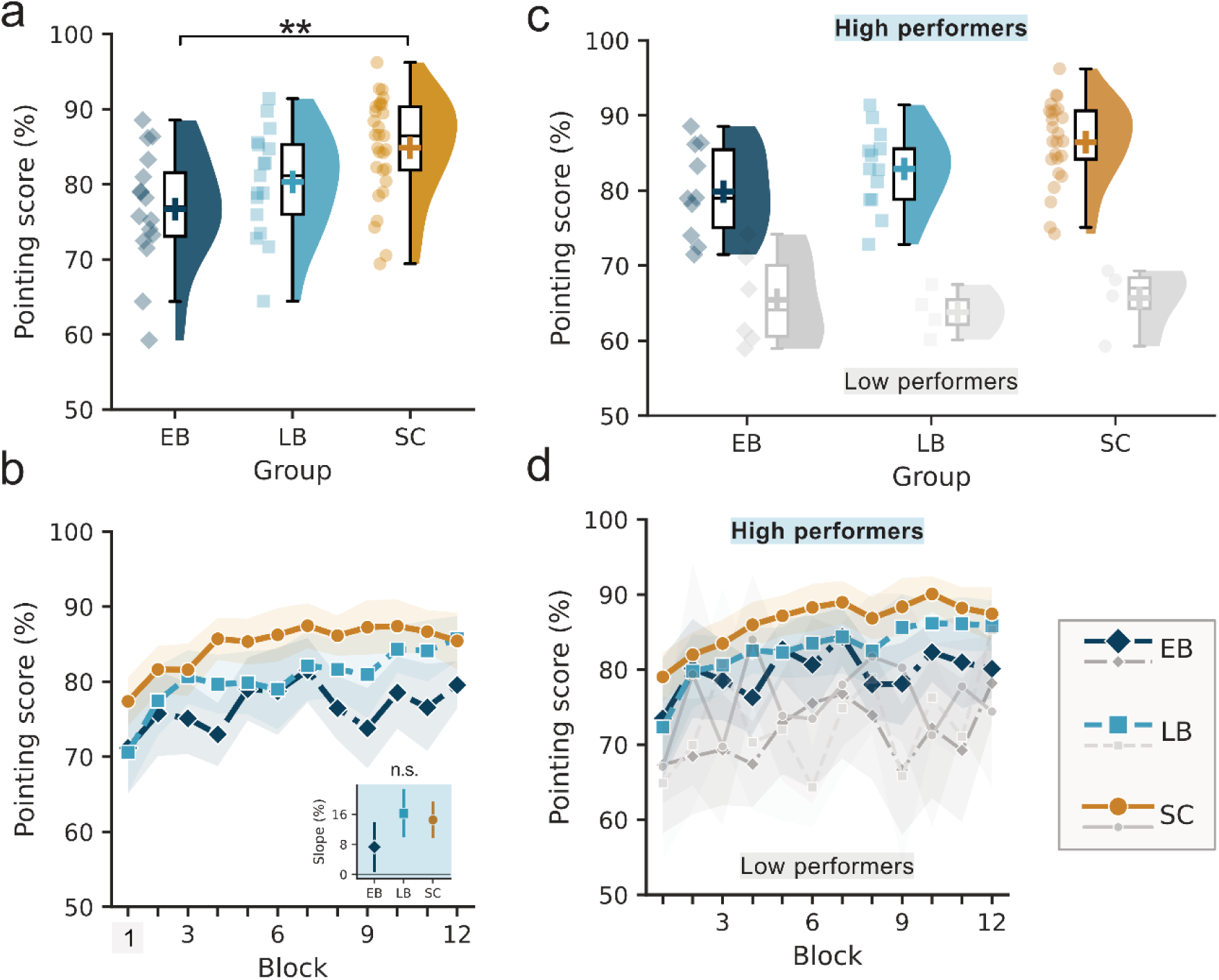
Average pointing scores and learning curves. **a.** Average pointing scores for each participant (individual data points) and their distribution; group averages are shown with crosses. **b.** Improvements in pointing scores across the 12 blocks; no differences in slope were detected. **c.** Average pointing scores for high (bold, colored) and low performers (thin, greyscale). **d.** Improvements in pointing scores across the 12 blocks for high (bold, colored) and low performers (thin, greyscale). For each error bar plot displaying learning rates, areas around group lines depict confidence intervals (95%). **, p<.01. Abbreviations: EB, Early Blind; LB, Late Blind; SC, Sighted Controls.

LME models of high- and low-performers’ pointing scores also best described the data using a power-law approach to asymptote (see SI Appendix). High performers did not show a difference at baseline (χ²(2) = 2.64, p = .267) nor a group × block interaction (χ²(2) = 4.13, p = .127), but the group difference in overall pointing accuracy was close to significance (χ²(2) = 5.96, p = .051). Exploratory pairwise contrasts showed that SC pointed more accurately than EB (β = 5.31, z = 2.43, p = .015, p_holm_ = .046) and LB were intermediate between SC and EB (p_holm_ = .227 and .354, respectively). Among low performers, no effect was detected (group × block χ²(2) = 0.24, p = .886; level χ²(2) = 1.32, p = .516; baseline χ²(2) = 0.31, p = .857) and no group improved reliably (see SI Appendix). Together, these results indicate that direction estimation is less accurate in EB, even in high-performing participants. More information is available in the SI Appendix.

### Mental Rotation (MR) abilities reveal a general blindness effect

MR abilities were examined as an additional measure of allocentric spatial cognition. Because participants learned the maze from a fixed viewpoint (facing North), trials requiring pointing from other orientations forced them to mentally reorient within the map (e.g., facing East/West: ±90° MR; facing South: 180° MR; see Fig. 4a). MR was analyzed with an LME model of pointing scores (3 MR levels × 3 groups) with age and sex as covariates (see Methods), and a random intercept per participant. Omnibus tests revealed a group × MR interaction (χ²(4) = 18.73, p < .001; see Fig. 4b), along with main effects of MR (χ²(2) = 30.75, p < .001) and group (χ²(2) = 11.94, p = .003). Furthermore, the groups did not differ at the 0° baseline level (Wald χ²(2) = 0.51, p = .774), suggesting group differences reflect differences in MR abilities rather than overall pointing abilities.

**Figure 4.**
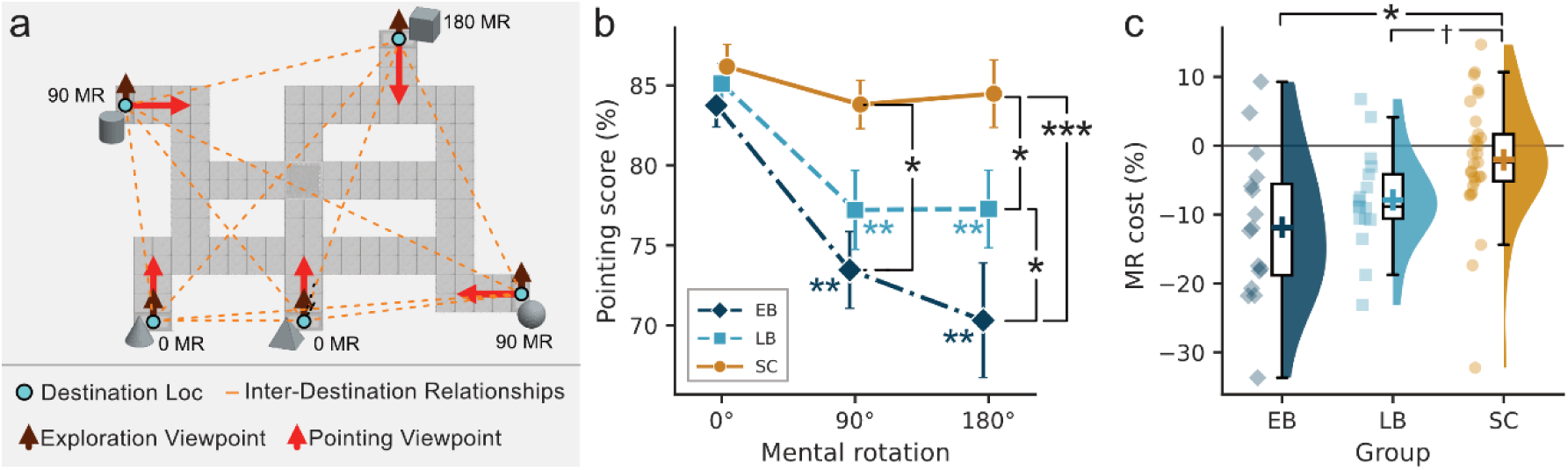
Mental rotation abilities. **a.** Maze layout, including the locations of all destinations (sphere, cube, pyramid, cone and cylinder), the exploration viewpoints (as experienced when learning the tactile maze), the viewpoints from each destination (used in the JRD pointing task), and the spatial relationships (or directions and distances) between all pairs of destinations (e.g., directions to compare with participants’ pointing data). **b.** Pointing scores declined significantly under mental rotation for EB and LB. **c.** Average MR cost for each participant (individual data points) and their distribution; group averages are shown with crosses. †, p<.10; *, p<.05; **, p<.01; ***, p<.001. Abbreviations: EB, Early Blind; LB, Late Blind; SC, Sighted Controls. Loc, Location; MR, mental rotation.

Pointing scores declined with increasing MR demand in both blind groups but not in SC. Relative to the 0° baseline, EB were less accurate at 90° (Δ = −10.28%, z = −4.66, p < .001, p_holm_ < .001) and at 180° (Δ = −13.43%, z = −6.09, p < .001, p_holm_ < .001), and LB showed the same pattern (90°: Δ = −7.90%, z = −3.69, p < .001, p_holm_ < .001; 180°: Δ = −7.83%, z = −3.66, p < .001, p_holm_ < .001), while SC were unaffected at either level (90°: Δ = −2.37%, z = −1.45, p = .148, p_holm_ = .444; 180°: Δ = −1.70%, z = −1.04, p = .300, p_holm_ = .601). At 90° MR, EB pointed less accurately than SC (Δ = −9.81%, z = −3.48, p < .001, p_holm_ = .002), with LB intermediate between SC and EB (p_holm_ = .096 and .182, respectively). At 180° MR, all groups differed: EB were less accurate than SC (Δ = −13.63%, z = −4.83, p < .001, p_holm_ < .001) and LB (Δ = −7.39%, z = −2.36, p = .018, p_holm_ = .037), and LB were less accurate than SC (Δ = −6.24%, z = −2.19, p = .029, p_holm_ = .037). No differences emerged between 90° and 180° for either group (all p_holm_ ≥ .153), suggesting MR costs did not scale with rotation size. Fig. 4b summarizes these effects, and the SI Appendix provides more detail.

Finally, MR was summarized for each participant with a single index: MR cost (in % of pointing score; mean(Score_90°_, Score_180°_) - Score_0°_). Overall, EB had an MR cost of −11.85 ± 11.01 %, LB, of −7.86 ± 7.27 %, and SC of −2.03 ± 9.18 % (see Fig. 4c). An ANCOVA with age and sex as covariates revealed a significant group effect (F(2,57) = 5.71, p = .005, η²_p_ = 0.167): relative to SC, MR costs were larger in both EB (t(25.9) = −2.83, p = .009, p_holm_ = .027, d = −0.91) and LB, to a lesser extent (t(40.7) = −2.09, p = .043, p_holm_ = .087, d = −0.62), while both blind group did not significantly differ (t(24.7) = −1.32, p = .200, p_holm_ = .200, d = −0.46). Taken together, these results suggest that MR costs are significantly higher in both blind groups, but most pronounced in EB.

### Pointing accuracy tends to increase with blindness duration

Overall, pointing accuracy (r = .79, p < .001) and MR cost (r = .50, p < .001) significantly correlated with navigation performance, controlling for age, sex, and group. When looking at these relationships by group, the pointing accuracy relationships held within each group, while MR was only significant in EB (see Table 2 and Fig. 5a). We also tested how age (corrected for sex) and duration of blindness (BDI, corrected for age and sex) are related to navigation performance, pace, pointing accuracy, and MR cost. Overall, SC older participants had worse navigation performance and pointing accuracy, while pace and MR cost did not significantly correlate with age. No age correlations were found in LB and EB. Still, correlation coefficients were positive in EB, contrary to SC and LB (see Table 2 and Fig. 5b). Comparing correlations across groups with Fisher r-to-z tests, the age-pointing correlation in EB was significantly more positive than in SC (r = +.41 vs r = −.52, z = 2.89, p = .004, p_FDR_ = .015). In contrast, the corresponding difference for navigation performance was not significant (EB vs SC, z = 1.83, p = .067, p_FDR_ = .134).

**Figure 5.**
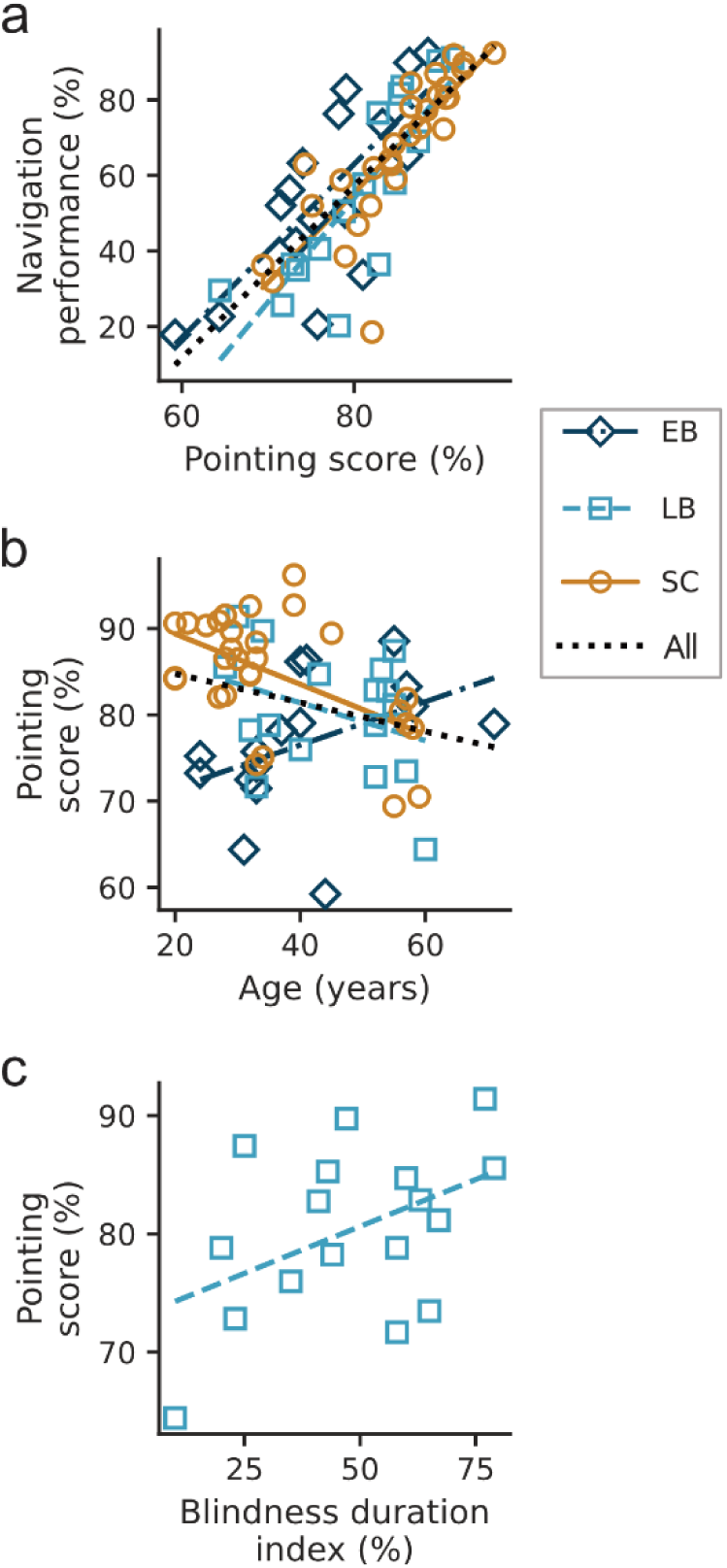
Correlation analysis of navigation performance, pointing scores, age, and blindness duration index (BDI). Scatter plots with trend lines show how **a.** average pointing scores relate to average navigation performance, **b.** age relates to pointing score, and **c.** BDI relates to pointing score. For each error bar plot displaying learning rates, areas around group lines depict confidence intervals (95%). **, p<.01. Abbreviations: EB, Early Blind; LB, Late Blind; SC, Sighted Controls.

While age was negatively associated with pointing accuracy in LB (though not significantly), an additional exploratory analysis revealed that blindness duration index (BDI, as % of life spent with blindness; see methods), when uncorrected for multiple tests, tended towards a positive association with pointing accuracy (r = .48, p = .071, p_FDR_ = .282; see Fig. 5c and SI Appendix). Taken together, this exploratory analysis suggests that aging mostly reduces pointing accuracy in SC, while this pattern is reversed in EB. Furthermore, correlations in EB and LB suggest that pointing accuracy tends to improve with blindness duration (in EB, BDI ≈ age). This qualitative observation warrants further testing with larger study samples.

Finally, we collected self-reported sense of direction via the Santa Barbara Sense of Direction (SBSOD) questionnaire and prior exposure to virtual environments via video game experience from blind participants (Table 1). SBSOD was associated (controlling for group, age and sex) with navigation performance (r = .52, p = .007, p_FDR_ = .014), pointing accuracy (r = .61, p = .001, p_FDR_ = .003) and MR cost (r = .43, p = .030, p_FDR_ = .040), but not with pace (r = −.38, p = .056, p_FDR_ = .056, see Table 2). Fourteen of 31 blind respondents had prior video game experience. Still, this proportion did not differ between EB and LB (8/16 versus 6/15, odds ratio = 1.50, p = 0.722) or between high and low performers (9/21 versus 5/10, odds ratio = 0.75, p = 1.000), and was unrelated to SBSOD (4.73 versus 5.10, Mann–Whitney U = 98, p = 0.793). Video game experience was also not related to any outcome (see SI Appendix).

**Table 1.**
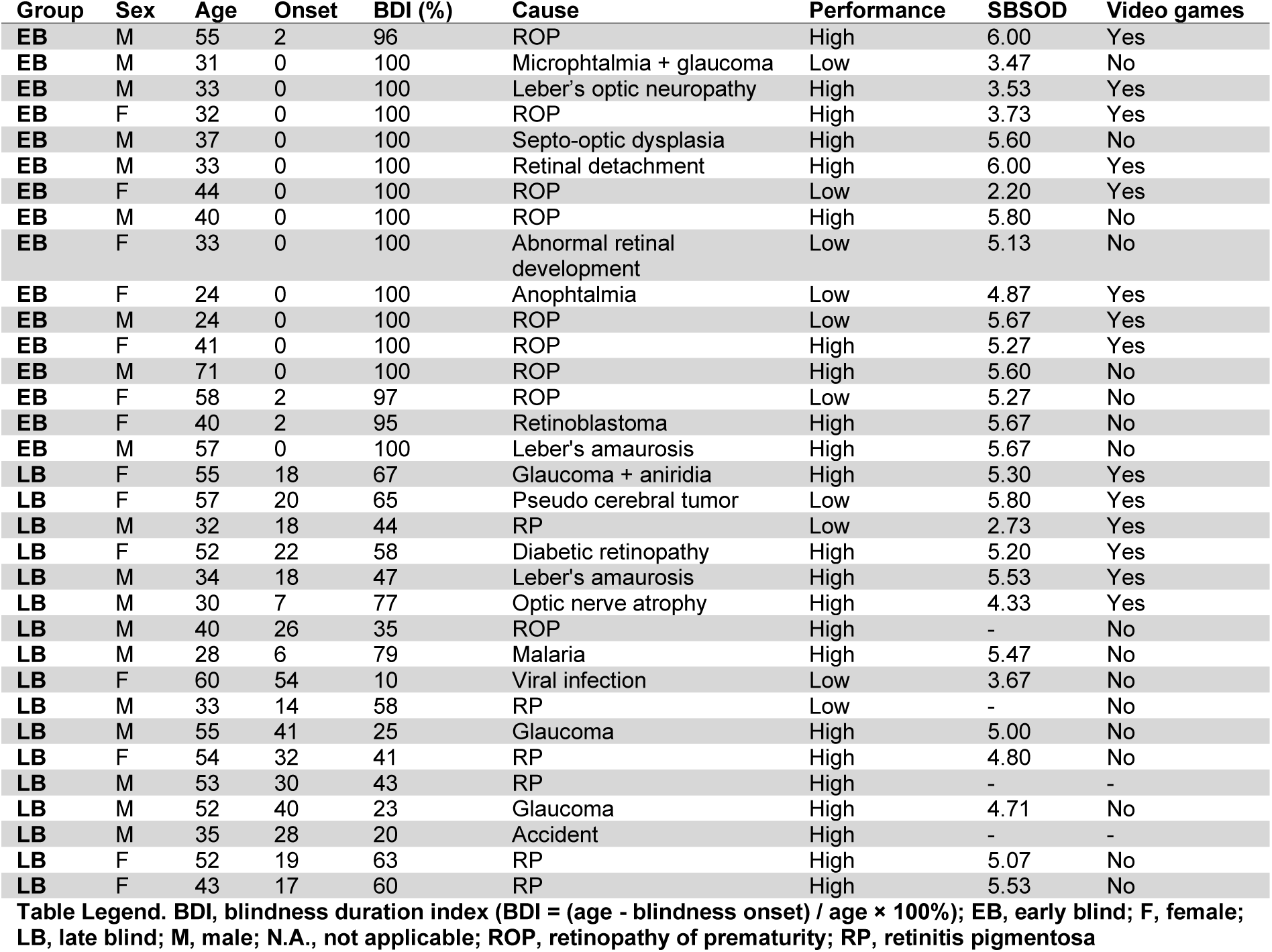
Characteristics of EB and LB participants.

| Group | Sex | Age | Onset | BDI (%) | Cause | Performance | SBSOD | Video games |
| --- | --- | --- | --- | --- | --- | --- | --- | --- |
| EB | M | 55 | 2 | 96 | ROP | High | 6.00 | Yes |
| EB | M | 31 | 0 | 100 | Microphthalmia + glaucoma | Low | 3.47 | No |
| EB | M | 33 | 0 | 100 | Leber's optic neuropathy | High | 3.53 | Yes |
| EB | F | 32 | 0 | 100 | ROP | High | 3.73 | Yes |
| EB | M | 37 | 0 | 100 | Septo-optic dysplasia | High | 5.60 | No |
| EB | M | 33 | 0 | 100 | Retinal detachment | High | 6.00 | Yes |
| EB | F | 44 | 0 | 100 | ROP | Low | 2.20 | Yes |
| EB | M | 40 | 0 | 100 | ROP | High | 5.80 | No |
| EB | F | 33 | 0 | 100 | Abnormal retinal development | Low | 5.13 | No |
| EB | F | 24 | 0 | 100 | Anophtalmia | Low | 4.87 | Yes |
| EB | M | 24 | 0 | 100 | ROP | Low | 5.67 | Yes |
| EB | F | 41 | 0 | 100 | ROP | High | 5.27 | Yes |
| EB | M | 71 | 0 | 100 | ROP | High | 5.60 | No |
| EB | F | 58 | 2 | 97 | ROP | Low | 5.27 | No |
| EB | F | 40 | 2 | 95 | Retinoblastoma | High | 5.67 | No |
| EB | M | 57 | 0 | 100 | Leber's amaurosis | High | 5.67 | No |
| LB | F | 55 | 18 | 67 | Glaucoma + aniridia | High | 5.30 | Yes |
| LB | F | 57 | 20 | 65 | Pseudo cerebral tumor | Low | 5.80 | Yes |
| LB | M | 32 | 18 | 44 | RP | Low | 2.73 | Yes |
| LB | F | 52 | 22 | 58 | Diabetic retinopathy | High | 5.20 | Yes |
| LB | M | 34 | 18 | 47 | Leber's amaurosis | High | 5.53 | Yes |
| LB | M | 30 | 7 | 77 | Optic nerve atrophy | High | 4.33 | Yes |
| LB | M | 40 | 26 | 35 | ROP | High | - | No |
| LB | M | 28 | 6 | 79 | Malaria | High | 5.47 | No |
| LB | F | 60 | 54 | 10 | Viral infection | Low | 3.67 | No |
| LB | M | 33 | 14 | 58 | RP | Low | - | No |
| LB | M | 55 | 41 | 25 | Glaucoma | High | 5.00 | No |
| LB | F | 54 | 32 | 41 | RP | High | 4.80 | No |
| LB | M | 53 | 30 | 43 | RP | High | - | - |
| LB | M | 52 | 40 | 23 | Glaucoma | High | 4.71 | No |
| LB | M | 35 | 28 | 20 | Accident | High | - | - |
| LB | F | 52 | 19 | 63 | RP | High | 5.07 | No |
| LB | F | 43 | 17 | 60 | RP | High | 5.53 | No |

Table Legend. BDI, blindness duration index ( $BDI = (age - blindness\ onset) / age \times 100\%$ ); EB, early blind; F, female; LB, late blind; M, male; N.A., not applicable; ROP, retinopathy of prematurity; RP, retinitis pigmentosa

**Table 2.**
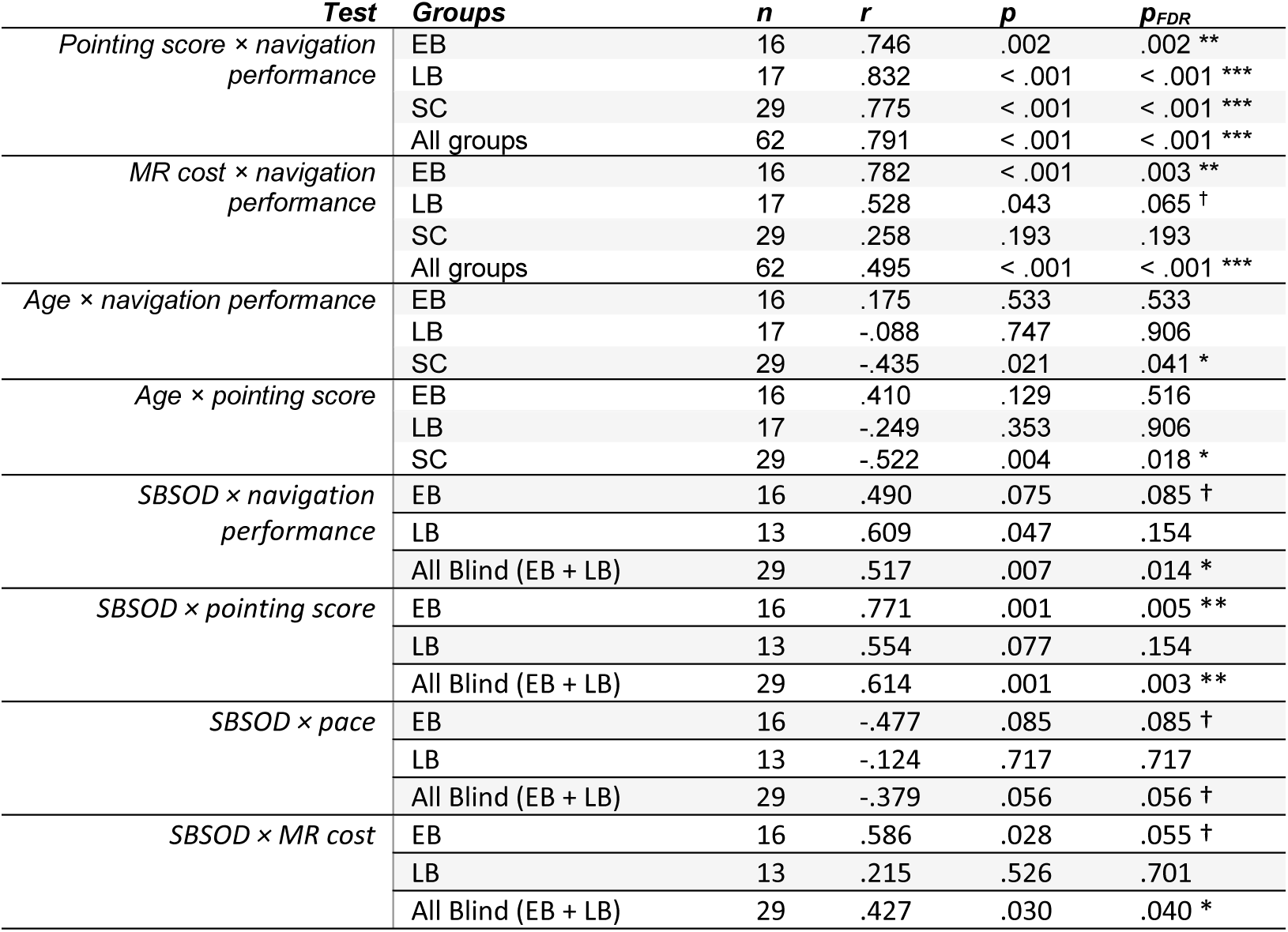
Partial Pearson results on the four outcomes (navigation performance, pace, pointing score, and mental rotation (MR) cost), age, and SBSOD.

## Discussion

In this study, we used an original “spatial learning” paradigm to monitor cognitive map formation in cases of early- and late-onset blindness. Most importantly, the results showed that EB, LB, and SC differ not only in allocentric pointing ability, but also in the temporal aspects of cognitive map formation, or, in other words, how fast they accumulate spatial knowledge for navigation. These results help clarify previous findings by indicating that differences between blind and sighted participants are not solely reflected in overall performance, but rather in how individuals acquire and use spatial knowledge over time. Our data further revealed substantial interindividual variability in navigation performance and pointing accuracy in all groups, which can be leveraged to identify and train behavioral and neural factors associated with higher spatial abilities in blindness.

### Blindness affects the temporal dimension of cognitive map formation

By testing whether EB, LB, and SC differ in learning rates, baseline performance, and overall performance, the study provides a quantitative framework for testing theoretical models of spatial knowledge acquisition in blindness (8, 35, 46, 60). Within this framework, equivalent initial navigation performance and slower learning rates argue against a persistent disadvantage and are consistent with cumulative or convergent models. Indeed, in these models, spatial knowledge is acquired successfully without visual experience but is accumulated and/or integrated more slowly into higher-order spatial representations (8, 35, 46). This distinction matters, as reporting group means alone would have misleadingly obscured this learning effect (7).

While this pattern is consistent with previous work showing that EB individuals have distortions in their spatial representations (4, 21–26, 28), it requires nuance. First, low performers were proportionally more frequent in EB (though not significantly), which could indicate, under the task constraints, differences in the time required to reach a given performance rather than ability to learn per se. Second, EB high performers showed no differences compared to SC and LB, suggesting that, depending on the individual, spatial knowledge can be acquired equally well. Third, EB performance also fit a quadratic function (see SI Appendix), consistent with a fatigue-like cost. This could reflect the less frequent exercise of the tested abilities, which are known to improve with training and access to spatial information (7, 21, 31, 36, 59, 64). Consistently, our findings support that more autonomous travelers (higher SBSOD) built better survey representations (62, 75, 76).

As a result, the present study further demonstrates that, when given access to survey knowledge via tactile maps, EB and LB individuals form cognitive maps, improve them, and navigate new spaces from memory, and they do so successfully despite differences in their mental representations (77). This holds even in virtual environments, despite the absence of self-motion cues (78, 79) and limited video game experience. We also measured faster navigation speeds in EB at baseline, which may reflect supranormal auditory processing during navigation, a mechanism that, while not specifically tested here, is well established in the literature (80–82). Furthermore, a subset of EB and LB participants reached expert map-use levels (high navigation and pointing scores), demonstrating that such high-level, allocentric spatial representations are attainable without visual experience (7, 35, 46, 59, 62, 64, 83, 84). This suggests experiential factors, rather than fundamental “deficits”, drive the observed variability in navigation strategies (85) and performance (29), and that apparent group differences could be reduced or eliminated through training or sensory augmentation (64, 86).

### Altered allocentric spatial cognition in EB and LB

From another perspective, navigation performance alone reflects how cognitive maps are used and linked to external cues during navigation, but not how they are internally constructed. Therefore, to assess cognitive map accuracy, we paired navigation with a pointing task, whose scores more closely reflected mental representations of spatial relationships (e.g., directions). The results showed that EB were less accurate than the other groups, supporting the idea of distorted mental representations of directions and distances (4, 21–26, 28), whereas visuo-spatial imagery in SC and LB may support more accurate representations (21, 29, 87, 88).

However, our findings further revealed that mental rotation disproportionately affected both blind groups, who showed reduced pointing consistency across viewpoints. The extent to which pointing accuracy declines under MR provides insight into the ability to flexibly switch between viewpoints, a process that imposes a higher cognitive load when using egocentric, route- or movement-based strategies (21, 25, 27, 59). Therefore, spatial disadvantages in EB and LB seem to stem from translating the maze’s survey viewpoint into egocentric “first-person” representations from different viewpoints. These challenges, arising during the transformation of spatial representations, may reflect differences in strategy rather than deficits in allocentric coding per se, consistent with what the literature has already proposed (4, 18, 21, 27, 59, 89–92).

On that note, while disadvantages often observed in EB (21, 25, 27, 59) could be attributed to the absence of visual experience and mental imagery (29), intermediate performance in LB, who had typical visual development, reinforces the idea that behavioral adaptations to blindness (not the absence of visual experience) seem detrimental to allocentric strategies (76, 93, 94), in favor of abstract cognitive mapping strategies (e.g., graph-based cognitive maps (95)) and route following. In other words, adapting to limited distal cues and sequential sensory integration from touch and audition (4, 96) can lead to underutilization of allocentric cognitive maps (4, 97–99) but does not prevent their development. Indeed, as SC exhibited reduced pointing accuracy with age, blind participants, particularly EB, showed improvement as blindness duration increased (or with age in EB). This likely reflects the influence of experience and supports both the convergent model of spatial knowledge acquisition (8, 35, 46, 60) and the idea that non-visual allocentric abilities are trainable. Nonetheless, this effect warrants further confirmation.

### Perspectives for future research and rehabilitation

Perceiving distal cues, such as landmark locations (or points of interest), is crucial for developing the allocentric reference frame and, thus, building allocentric cognitive maps (1, 100, 101), but access to such cues remains one of the main challenges for individuals with blindness, despite available assistive technologies (102). Research, technology development, education, and urban planning must improve access to distal cues and facilitate survey viewpoints to enhance allocentric cognitive mapping in blindness (12, 14–16, 31, 62, 74, 84, 103, 104). For this purpose, multisensory virtual environments, such as the one used in this study, offer a promising avenue for providing: 1) tools to assess cognitive map formation independently of visual status and to target skills for reinforcement through Orientation & Mobility training (36, 105); and 2) multisensory games that progressively reinforce crucial but underutilized allocentric cognitive abilities, such as spatial inference (directions and distances) and perspective switching. Indeed, video game training has been shown to improve spatial abilities in blind children and in other clinical and non-clinical populations (106–109), yet access to such tools remains limited.

Finally, this study provides evidence that a virtual multisensory maze-learning task that combines pointing and navigation can yield a detailed understanding of cognitive maps and their formation over time. This spatial learning paradigm has strong adaptive potential for neuroimaging applications and for investigating spatial cognition in other clinical populations. Indeed, this fully automated task has low motor demands and is independent of participants’ language abilities, making it a potentially relevant investigative and training tool in the context of Alzheimer’s disease, Parkinson’s disease, traumatic brain injuries, aphasia, and many other conditions.

### Limitations

In this study, we quantified and tracked, over time, two aspects of cognitive map formation: a functional aspect, whether participants could reach remembered destinations, and a spatio-cognitive aspect, whether directions between locations were accurately represented (JRD task). Both improved and were strongly correlated, indicating that more accurate Euclidean relationships were linked to more successful wayfinding. This convergence does not, however, identify the format of the underlying representation, since the maze layout and the task would equally support metric and graph-like cognitive maps (95), nor does it establish which strategy individual participants adopted. Pairing these measures with a final map reconstruction task, which this protocol did not include, would have been an effective way to probe the structure of the underlying cognitive map. The virtual navigation task in the present study also does not directly reflect real-world navigation abilities. Indeed, static virtual tasks do not provide vestibular and proprioceptive feedback, which are integral to real-world navigation (78, 79). In sighted participants, these body-based cues improve spatial updating and cognitive mapping (110–112), and their absence may weigh more heavily in blindness, although this has not yet been directly tested. Not all participants were familiar with virtual environments, and some blind participants therefore adopted conservative strategies (i.e., slower decision-making to avoid errors). We believe these adaptive strategies should not be interpreted as deficits, as blind individuals must prioritize safety and certainty over speed in daily travel. Finally, the study involved only one controlled virtual environment, limiting generalizability. However, the fact that spatial knowledge from similar virtual tasks translates to real-world settings (32, 66–71) provides strong confidence in the validity of the present findings.

## Conclusion

The present study offers a scalable, quantitative framework for monitoring the formation of non-visual cognitive maps over time through virtual navigation and pointing. It also improves ecological validity in the context of blindness by focusing on active exploration and sequential information integration, rather than visual guidance (113). As all groups improved in navigation and pointing, findings reinforce that cognitive maps are fundamentally cross-modal constructs that do not depend on visual experience and are highly trainable in both EB and LB. However, the results revealed that differences between blind and sighted individuals are primarily in the temporal dimension of cognitive map formation, with EB individuals accumulating spatial knowledge for navigation more slowly. Furthermore, reduced pointing accuracy under viewpoint transformation in EB and LB, but not SC, suggests that blindness changes cognitive mapping strategies, even with considerable visual experience. Importantly, this emphasizes that cognitive map formation is an experience-dependent process that varies across time and individuals, both dimensions that have received little attention in the literature.

## Materials and Methods

### Participants and consent

A total of 62 participants took part in the study, including 33 with total blindness. These participants were classified into three groups: EB (n = 16, 9M, 7F, mean age = 40.8 ± 13.2 years), LB (n = 17, 10M, 7F, mean age = 45.0 ± 11.0 years), and blindfolded sighted controls (SC; n=29, 15M, 14F, mean age = 35.4 ± 12.5 years). For each blind participant, a blindness duration index (114) was calculated (BDI = (age - blindness onset) / age × 100%). Blind participants (16 EB, 13 LB; 4 LB could not be reached) also completed the Santa Barbara Sense of Direction (SBSOD) scale questionnaire (115, 116), which assessed their general sense of direction and independence, and reported whether they had experience with video games (including both mainstream and specialized audio-games). Table 1 provides additional information on EB and LB participants.

All participants provided informed consent before participating. The study was conducted between 08/01/2023 and 04/01/2026 in accordance with the Declaration of Helsinki and approved by three ethical committees (CERC of the University of Montreal, Montreal, Canada; CERVN of the CCSMTL, Montreal, Canada; Region H committee, Copenhagen, Denmark). We recruited all EB and LB participants from specialized organizations and institutions for the visually impaired, and through snowballing; we recruited SC participants through public digital invitations and snowballing. The behavioral experiment was conducted on-site in French or English (according to the participant’s preference). Questionnaires, available in the SI appendix, were administered on-site or by phone after the experiment (if time constraints required it). All participants followed the same training procedures before proceeding to the experiment (see SI appendix).

### Material

#### Maze design

The maze represented a complex environment: a maze-like network of interconnected paths with multiple destinations. Destinations were represented by five 3D geometric shapes (sphere, cube, pyramid, cylinder, and cone). This design was chosen to prevent participants from learning specific routes and to encourage a cognitive map-based strategy that allows them to improvise their way from one random destination to another. This multiple route design made it difficult to memorize all routes within the allocated exploration time (60s; Experimental procedures). Indeed, this system allowed for 20 different routes, of which 16 were presented.

Destination placement followed the maze’s two axes: four destinations occupied opposite quadrants (Sphere = South-East, Cube = North-East, Cylinder = North-West, Cone = South-West), and the fifth (Pyramid) was placed centrally on the southern side and was used as the reference point where the experimenter placed the participant’s index at the start of the first exploration phase. Trials were constructed so that every route required crossing the environment along at least one axis (North-South or East-West). Because the pyramid was central, the *Sphere-Pyramid* and *Pyramid-Cylinder* pair routes were not presented. The maze distance unit was a “tile”, echoing the tile system of the virtual environment (participants could move from tile to tile; see the *Audio-based Virtual Environment* section), and all hallways were two tiles wide. On average, the mean shortest-path length was 23.03 ± 4.55 tiles (range = 16-32 tiles), the mean Euclidean distance between destinations was 16.91 ± 3.51 tiles (range = 11.60-23.26 tiles), and the number of minimal turns per route was 3.29 ± 0.77 (range = 2-5 turns). Finally, route circuitousness was indexed by the detour ratio, or the shortest walkable path divided by the Euclidean distance between the two destinations. Presented routes had a mean detour ratio of 1.39 ± 0.13 (range = 1.23-1.68), so the shortest possible route was on average 39% longer than the Euclidean, direct line. Table 3 shows route information for each location pair.

**Table 3.**
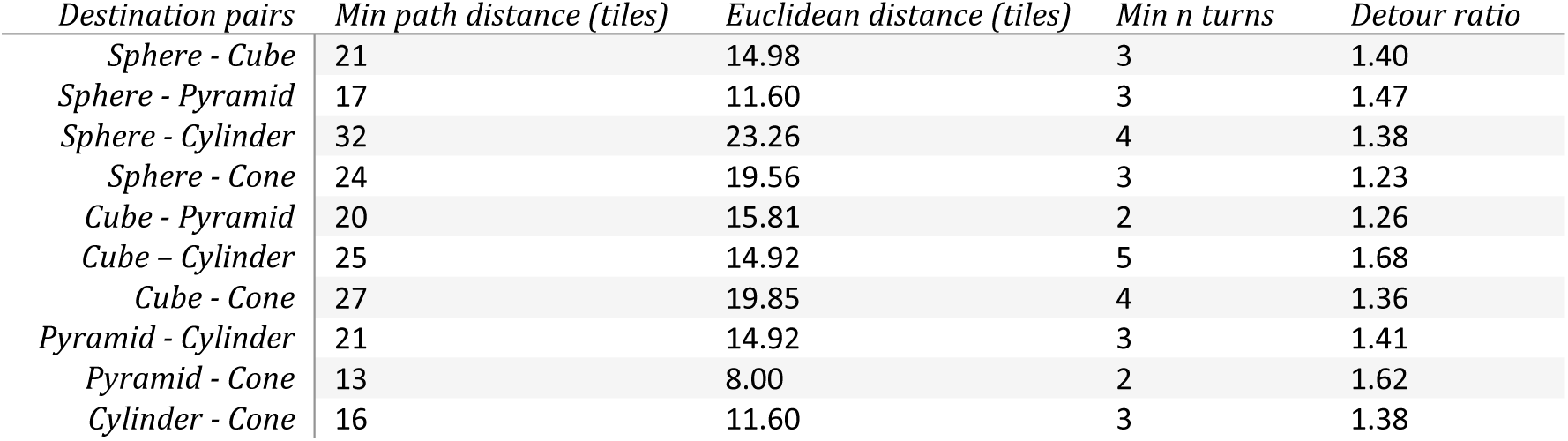
Route information for each destination pair.

| Destination pairs | Min path distance (tiles) | Euclidean distance (tiles) | Min n turns | Detour ratio |
| --- | --- | --- | --- | --- |
| Sphere - Cube | 21 | 14.98 | 3 | 1.40 |
| Sphere - Pyramid | 17 | 11.60 | 3 | 1.47 |
| Sphere - Cylinder | 32 | 23.26 | 4 | 1.38 |
| Sphere - Cone | 24 | 19.56 | 3 | 1.23 |
| Cube - Pyramid | 20 | 15.81 | 2 | 1.26 |
| Cube - Cylinder | 25 | 14.92 | 5 | 1.68 |
| Cube - Cone | 27 | 19.85 | 4 | 1.36 |
| Pyramid - Cylinder | 21 | 14.92 | 3 | 1.41 |
| Pyramid - Cone | 13 | 8.00 | 2 | 1.62 |
| Cylinder - Cone | 16 | 11.60 | 3 | 1.38 |

#### 3D printed tactile maze

To develop their cognitive map of the complex environment, participants were given a tactile board with a 15×10 cm physical rendition of the previously described maze; it comprised a 3D-printed set of hallways and destination shapes (Fig. 1a). To learn the maze, participants had to freely explore it with their dominant hand (Fig. 1b). This aspect of the task was adapted from a previous protocol (62).

#### Audio-based Virtual Environment

To quantitatively assess the content and precision of participants’ cognitive maps, we used an audiovisual game inspired by previous studies (32, 51, 66) that we developed in-house using JavaScript, Python, and the tile map editor from the RMMV game engine (117). To perform the task, participants received auditory instructions and sounds via headphones and interacted with the software using a customized keypad. The keypad had nine usable buttons, including five with differently oriented bars (from −90 to 90 degrees, in 45-degree increments; for the pointing task), three arrow keys (up, left, right; for virtual movements during the navigation task), and an additional button (symbol: two vertical bars; used to perform a side-step). The keypad also had a central tactile marker where participants placed their index fingers between tasks. Fig. 1a shows the entire apparatus. A complete description of the virtual environment and related hardware is available in the SI Appendix, and movie S1 shows how participants experienced it.

### Experimental procedures

We used an original “spatial learning” paradigm designed to track maze learning (cognitive map formation) across repeated periods of free exploration and virtual navigation. We assessed learning progress at several time points by measuring participants’ performance in the virtual, life-size maze. The paradigm was organized into 12 experimental blocks, each comprising three phases that measured different aspects of cognitive map-based navigation (Fig. 1a; see Video S1):

1. **Exploration (or learning) phase (60 s).** Participants freely explored the tactile maze (using the index finger of the dominant hand) to learn the map as much as possible (cognitive map formation). In this phase, participants were explicitly tasked with learning the overall layout of the maze (not individual routes) and the relative positions and directions (see SI Appendix). A 6-second pause then followed.
2. **Pointing phase (12 s).** This phase tested cognitive map retrieval and anchoring across different locations and viewpoints. The audio-based game first announced a random pair of start location and destination: “*Start: [start location]; Goal: [destination].*” Then, the participant was asked to point (instruction: “*point*”) toward the destination (relative to the specific viewpoint of the start location; see Fig. 4a). This constituted a judgment of relative direction (JRD) probing the allocentric relationships represented in the cognitive map. The facing direction (or the locations’ viewpoint) was not verbally given; participants had to retrieve it from memory based on the maze geometry. Participants then entered their response by pressing and holding their direction estimate for the remainder of the pointing phase (6 s). Participants had 9 possible answers (from −90 to 90 degrees, in 22.5-degree increments), entered by pressing one or two buttons at a time. A 6-s pause followed.
3. **Navigation phase (maximum 25 s).** To test how participants used cognitive maps for navigation, they navigated the audio-based virtual environment from the start location to the destination using the keypad (arrow keys; Fig. 1a). The 25-second ceiling was set at twice the fastest time to complete the longest route (sphere to cylinder; 12.5 s). It was applied for three reasons: 1) motivating participants to learn the general layout (not specific routes) and reach destinations as efficiently as possible; 2) stopping lost participants from endlessly wandering in the maze; and 3) constraining participants to the task (reaching the destination as efficiently as possible), while keeping them from learning the maze during virtual navigation (e.g., virtual exploration). A 6-s intertrial interval (ITI) followed, after which a new trial began with a new pointing phase.

In sum, the experiment consisted of 12 experimental blocks, each with one exploration phase and 8 trials, each consisting of one pointing phase and one navigation phase. All participants completed the entire experiment except for 8 participants (3 EB, 2 LB, and 3 SC) who could not complete it due to time constraints. This resulted in 5835 collected trials for analysis. All participants were included in the analysis.

### Behavioral markers of cognitive map formation

#### Navigation and Pointing

For statistical analysis, multiple metrics were collected for each trial during the pointing and navigation tasks. In the navigation task, three variables were collected in each trial: 1) *Navigation success or failure*, depending on whether the correct destination was reached; 2) *Time-to-destination*, or the time (in seconds) needed to reach a destination in both successful and failed trials (trials ended due to the 25s ceiling were censored); and 3) the “*End position”, or the* participant’s x and y coordinates (in tiles) at the moment the trial ended. These three variables were used to define *Navigation Performance* (%) and *Pace* (s per tile) to monitor participants’ improvement throughout the experiment, per trial and per block (8 trials per block).

In each trial, navigation performance was quantified as follows: 100% (Success, destination was reached); 90% (end position was within a 3-tile radius of the destination); 75% (end position was within a 5-tile radius of the destination); 50% (end position was within a 7-tile radius of the destination), or 0% (failure, end position was beyond a 7-tile radius of the destination or the wrong destination was entered). This quantization avoided binary per-trial performance and made the average more representative of participants’ abilities. Indeed, participants may have been successfully navigating toward the correct destination but did not have time to reach it within the allocated 25 seconds. This quantization system was selected over distance measures because the task is non-visual: distance information does not account for situations in which participants are lost or randomly wandering in the maze, which would inflate performance (see SI Appendix). Thus, quantization ensured that percentage points were given only when the participant was approaching the correct destination.

For each trial, a pace measure (in seconds per tile) was computed and defined as the time to reach the destination divided by the straight-line (Euclidean) distance between the start and destination locations (see Table 3). Therefore, pace is defined only on trials that ended at the requested destination (3,054 of 5,835 navigation trials), and lower pace values indicate faster, more efficient travel. Because pace is computed only for successful trials, we fitted its models using weighted least squares with inverse-probability weights and clustered standard errors by participant; the three group comparisons are then joint Wald tests on that cluster-robust covariance (see SI Appendix).

During the pointing task, three variables were collected: 1) whether participants “*pointed*” by providing a direction estimate or not (*Boolean:* 0 = not pointed, 1 = pointed); 2) the *direction estimate* (participants’ answers in degrees, between −90 and 90 degrees, 0 degrees = directly in front of the participant); and 3) an *angle delta* (Δ*θ*), defined as the difference between the direction estimate and the real destination direction (when accounting for the starting location viewpoint). With these three metrics, a normalized pointing score measure was defined for trials where participants provided a direction estimate (see Equation 1):

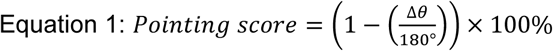

Furthermore, trials were divided into three categories based on how much the viewpoint from the starting location differed from the viewpoint used to experience the tactile maze (facing North). This categorization aimed to assess the cost of mental rotation (MR) on pointing scores and, therefore, participants’ allocentric coding abilities. Under this categorization, 0-degree MR trials started facing North (0-degree difference), 90-degree MR trials started facing East or West (90-degree difference), and 180-degree MR trials started facing South (180-degree difference). Finally, MR was summarised for each participant as a single index: MR cost (in % of pointing score; mean(Score_90°_, Score_180°_) - Score_0°_).

### High and low performer classification

To test whether group differences reflect genuine differences in how the maze is learned, rather than differences in how many participants failed or succeeded in learning it, participants were classified as high or low performers based on whether they learned the maze. This criterion was set at a threshold of 75% in at least one block, corresponding to six successful trials out of eight (6/8). With five possible destinations, a participant navigating without knowledge of the layout (and reaching a destination 100% of the time) would reach the requested destination in 1.6 of 8 trials (chance level). The criterion therefore represented the complement of chance (6.4 of 8) but was rounded down to 6 of 8, which accommodated trials in which no destination was reached.

### Statistical analyses

All statistical analyses were conducted with Python 3.11 (using *numpy, scipy, statsmodels, pandas*) and included linear mixed-effects (LME) models to analyze group differences in learning rates and overall performance.

#### Learning-curve form

Before conducting the LME model analyses, the form of the participants’ learning curve was empirically selected. Six candidate forms were tested: linear, logarithmic, power, exponential, quadratic, and block as an unordered factor. We fitted these forms with participants’ random effects and covariates (age and sex). We compared their sensitivity to the data using the Akaike information criterion, corrected for small sample sizes (AICc). These tests, and how they were conducted, are reported in the SI Appendix, and the most fitting form is stated for every main test conducted and reported in the main manuscript.

#### Linear mixed-effect models and omnibus tests

Maximum likelihood LME models were fitted to participants’ performance across all trials as input (trial-level data). For each variable tested, LME models were fitted according to the winning form (determined by the AICc) for this specific variable, separately for the whole sample, high performers, and low performers. Models included a random intercept per participant and, where the selected form contributes a single term, a random slope, as well as age and sex as covariates. The SI Appendix provides more information on the models’ equations. Then groups’ learning curves were compared in three different ways: (i) Slope (group × block interaction): does one group improve faster than another? (ii) Group main effect: is one group’s curve overall higher relative to another? (iii) Baseline: do the groups differ at block #1?

First, to test the group × block interaction, we fitted the LME model twice: once allowing each group its own slope (interaction model), and once sharing a single slope across groups. Then, a likelihood-ratio (χ²) test evaluated whether allowing groups to differ improved the LME fit significantly, indicating whether groups’ learning rates differed. Second, to test the main group effect, the same test was employed, but with a model in which each group has its own height (additive model) against one where they don’t. Third, to test baseline differences, we coded the block term so block #1 equals zero, and a Wald test (Wald χ²) evaluated whether the group coefficients in the first block differed significantly.

To describe these effects, or their absence, we report within-group slopes, learning-rate contrasts, and baseline contrasts from the interaction model, and overall group differences from the additive model. To obtain all three pairwise comparisons, each model was reparametrized with each group in turn as the reference level, giving identical predictions but placing different sets of comparisons in the coefficient vector. Beta coefficients (β) with z-scores (z = β/ standard error) and p-values (with and without Holm correction) are reported in the text. We report the complete outputs of each test in the SI Appendix.

#### Correlational analyses

We assessed relationships between the spatio-cognitive measures (pointing score and MR cost) and navigation performance using partial Pearson correlations, controlling for group, age, and sex; within each group, we controlled for age and sex. We also ran partial Pearson correlations to assess whether age (EB, LB, and SC; controlling for sex) and the BDI (LB only; controlling for age and sex) were related to each of the four outcomes (navigation performance, pointing scores, pace, and MR cost). We evaluated between-group differences in the age correlations using Fisher r-to-z tests on these same sex-adjusted correlations. Because these tests were exploratory, we applied Benjamini-Hochberg false discovery rate (FDR) correction within each family of tests. For associations between pointing scores and MR cost and navigation performance, we corrected correlations for three tests (EB, LB, and SC) each, and the two pooled estimates formed a separate family of two. For the remaining correlation analyses, each family comprised the four outcomes within a single group (correlations with age and BDI), and we corrected Fisher r-to-z tests across the four outcomes within each of the three group pairs (EB vs LB, EB vs SC and LB vs SC).

Blind participants’ self-reported sense of direction (SBSOD score) and reported video game experience were also each related to the four outcomes (navigation performance, pointing score, pace, and MR cost) by partial Pearson correlations, controlling for age and sex within EB and within LB, and for group, age and sex in the pooled blind sample. Following the same rule, we applied FDR correction across the four outcomes within each sample (EB, LB, and the pooled blind sample), separately for each of the two measures. Since the video-game item was binary, its coefficient reflects a partial point-biserial correlation. Finally, we tested the distribution of video game experience between groups (EB vs LB) and between high and low performers with Fisher’s exact test. We used a Mann-Whitney U test to evaluate whether participants differed in SBSOD score by video game experience.

## Acknowledgments

The team would like to thank all participants for their time and efforts.

## Data Availability

Data used in the study are available in the DRYAD repository ([DOI]). The code used to generate the results is available in the GitHub repository ([URL]).

