## Supplementary material for "Tracking acquisition of cognitive maps in early- and late-onset blindness": SI Appendix and Movie S1: REVISIONS_BIORXIV_SI-Appendix.pdf

**This PDF file includes:**

Legend for Movie S1
Supporting texts S1-S9
Figures S1-S7
Tables S1-S14

**Other supporting materials for this manuscript include the following:**

Movie S1

**Legend for Movie S1 (separate file).**

The movie follows the time course of the experimental procedures along one repetition of each experimental phase: one exploration phase, one pointing phase, and one navigation phase. This series would be followed by seven other trials consisting of one pointing phase and one navigation phase before reverting to the exploration phase.

Timings have also been adjusted for the purpose of minimizing the length of the video.

The video depicts what is seen by the experimenter on the computer monitor, and what is heard by the participants (all participants are blindfolded and do not see the monitor). At the bottom left, the video displays the phase number and name, and, at the bottom right, the apparatus (placed on participants' hips) is displayed.

Over the apparatus, a hand appears at different moments, identifying the times where participants interact with the apparatus (either exploring the tactile maze in phase 1, entering a direction estimation in phase 2, or moving around in the virtual maze using the arrow keys).

The experimental procedures follow the following descriptions:

1. Learning phase (60 s). Participants freely explored the tactile or visual maze (using the index finger of the dominant hand) to learn the map as much as possible (cognitive map formation). A 6-second pause then followed.

2. Pointing phase (12 s). This phase tested cognitive map retrieval and anchoring across different locations and viewpoints. The audio-based game first announced a random pair of start location and destination: "*Start: [start location]; Goal: [destination].*" Then, the participant was asked to point (instruction: "point") in the direction of the destination (as viewed from the start location). They entered their response by pressing and holding their direction estimate for the remainder of the pointing phase (6 s). Participants had 9 possible answers (from -90 to 90 degrees, in 22.5-degree increments), entered by pressing one or two buttons at a time. A 6-s pause followed.

**3.** Navigation phase (maximum 25 s). To test how cognitive maps are used for navigation, participants navigated the audio-based virtual environment from the start location to the destination using the keypad (arrow keys; Fig. 1a). A 6-s intertrial interval (ITI) followed, after which a new trial began with a new pointing phase.

#### Text S1 – The virtual environment

The virtual maze reproduced the layout of the 3D-printed tactile map at life size. It is a network of interconnected corridors on a square tile grid with five destinations (3D geometric shapes: sphere, cube, pyramid, cylinder and cone), hidden behind doors. The greatest distance (in tiles) between any two destinations constituted the maze diameter (23.26 tiles), a constant used to normalize distance-based scores (continuous spatial scoring, see text S2).

During the navigation phase, participants move with the arrow keys in discrete steps along the corridor grid. Participants have three different movement options: “walk forward one tile”, “turn 90 degrees to the [left or right]”, and a “sidestep” (Fig. S1a). If they walk forward, they hear a footstep sound effect and, if they turn, they hear a “whoosh” sound effect in the ear corresponding to the side they turned, followed by a verbal cue announcing the new cardinal direction they are now facing. The sidestep also triggers a distinct sound effect to announce the sidestep was successfully performed, it can only be performed if a wall is directly to the right or left of the participant.

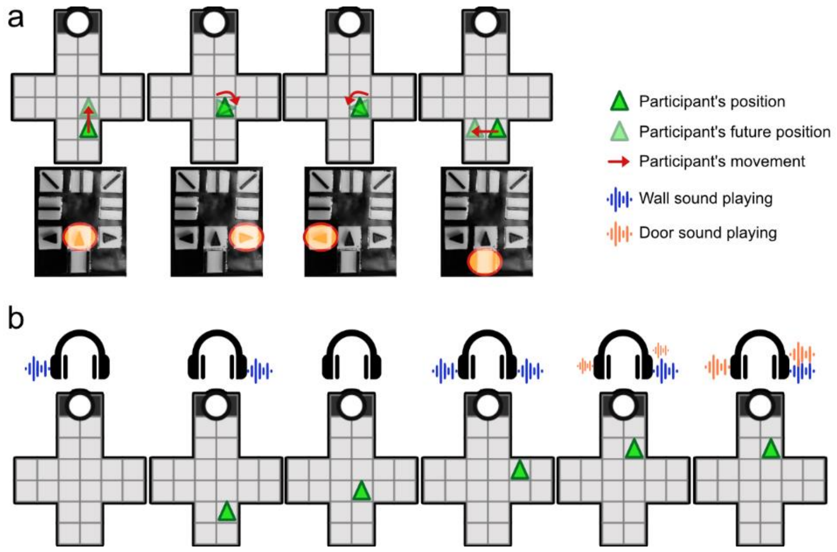


**Figure S1. The virtual environment.** A. The movements participants can perform and the buttons to perform them. B. Wall and door sound effects and how they are heard according to the participants’ position.

According to this system, participants can navigate in the hallways while hearing the walls and doors. Walls emit a looping sound effect that emulates a cane tapping on a wall and the doors emit a looping “door closing” sound effect. Walls are heard when the participant stands next to it and doors emit the sound in a two-tile radius and are louder when the participant is standing in the tile directly in front of the door. Figure S1b displays how the sounds are presented to the participants during navigation, Movie S1 shows one example of a successful navigation trial, and examples of the sound effects are available <u>here</u>.

**Text S2 – Navigation performance quantization**

In each trial, navigation performance was quantified as follows: 100% (Success, destination was reached); 90% (end position was within a 3-tile radius of the destination); 75% (end position was within a 5-tile radius of the destination); 50% (end position was within a 7-tile radius of the destination), or 0% (failure, end position was beyond a 7-tile radius of the destination or the wrong destination was entered). Figure S2 displays this system.

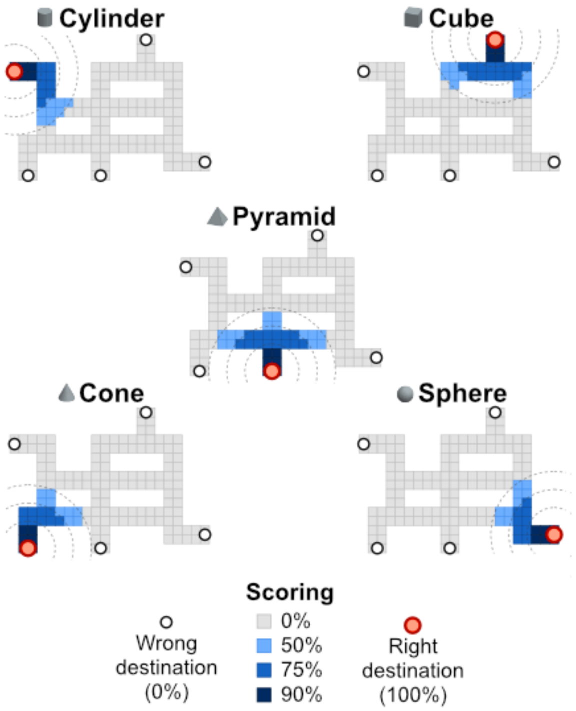


**Figure S2. Quantization of navigation performance according to the participant's final x and y position.** The virtual maze layout is displayed and colored according to trial destination (sphere, cube, pyramid, cone, or cylinder). Colors represent the navigation performance given to participants according to their X, Y position (tile position) at the end of the trial, relative to the destination. If participants reached a destination, we recorded a performance of 0 or 100%, depending on whether it was the correct destination.

This quantization avoided binary per-trial performance and ensured that percentage points were given only when the participant was approaching the correct destination. Indeed, participants may have been successfully navigating toward the correct destination but did not have time to reach it within the allocated 25 seconds.

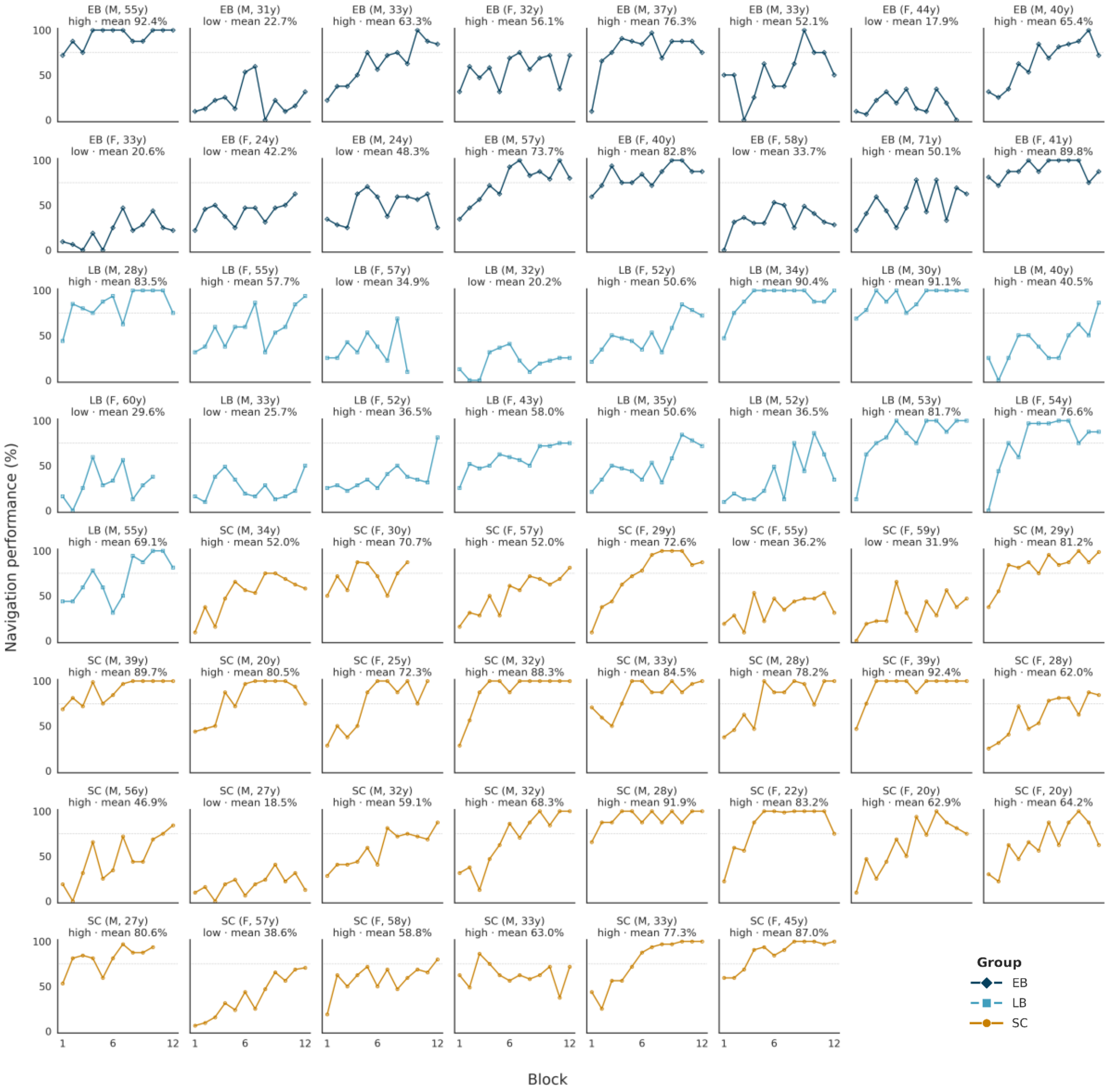


**Figure S3. Per-participant learning curves following the quantization system.** Each panel is one participant, labelled by group, sex and age, with their high/low performer classification and mean navigation performance over all their trials. Panels are ordered by group (EB, LB, SC) and each panel displays a dotted line marking the 75% criterion used to classify high and low performers.

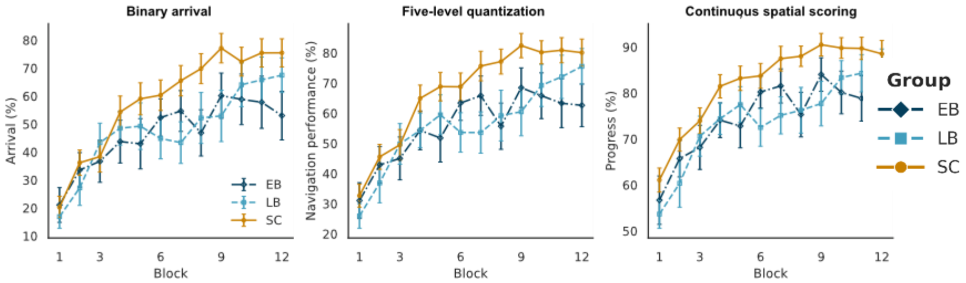


**Figure S4. Navigation performance, learning curves, scored three ways.** From left to right: 1) binary arrival (success or failure); 2) the five-level quantization used in the paper; and 3) continuous spatial scoring, or [100% × (1 − distance / maze diameter)] where distance represents the vector length (in tiles) between the destination coordinates and the participant's final coordinates. Points are group means of participant means, error bars are standard errors.

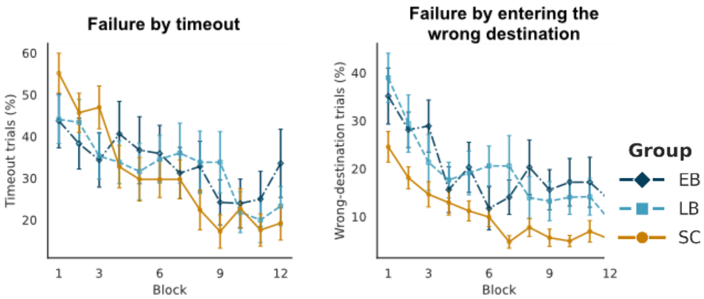


**Figure S5. Failed trials.** Proportion of timeouts and wrong destination trials over blocks, in % of failed trials.

The quantization system was selected over the continuous spatial scoring (distance measures) because distance information does not account for situations in which participants are lost or randomly wandering in the maze, which can be frequent due to the non-visual nature of the task (overestimate performance in failed trials). As possible to see in Figure S4, the continuous spatial scoring inflate performance compared to the binary arrival and five-level quantization scoring systems. Furthermore, the difference between baseline values and the maximum block performance is reduced in the continuous spatial scoring system. The omnibus tests on the LME models for the three systems are compared in Table S1; it is possible to see the continuous spatial scoring is not sensitive to group differences.

**Table S1. Omnibus tests under each scoring system, whole sample.**

| *Outcome* | *Form* | *Omnibus test* | *Test* | *$\chi^2$* | *df* | *p* |
|---|---|---|---|---|---|---|
| *Binary arrival* | exponential | Group x Block | LRT | 10.21 | 2 | .006 * |
| *Binary arrival* | exponential | Group | LRT | 0.12 | 2 | .942 |
| *Binary arrival* | exponential | Group at baseline | Wald | 0.45 | 2 | .797 |
| *5-Level quantization* | exponential | Group x Block | LRT | 8.77 | 2 | .012 * |
| *5-Level quantization* | exponential | Group | LRT | 1.13 | 2 | .568 |
| *5-Level quantization* | exponential | Group at baseline | Wald | 0.25 | 2 | .881 |
| *Continuous spatial scoring* | power | Group x Block | LRT | 2.24 | 2 | .327 |
| *Continuous spatial scoring* | power | Group | LRT | 3.71 | 2 | .156 |
| *Continuous spatial scoring* | power | Group at baseline | Wald | 1.33 | 2 | .516 |

#### Text S3 – Pointing task, judgment of relative direction (JRD)

For the pointing task, participants had to perform a JRD with the custom keypad (5 buttons depicting different directions from -90 degrees to +90 degrees; 9 possible answers by combining buttons; Fig. S6a) by imagining the maze as viewed from the viewpoint of the starting location (Fig. S6b-c), from memory; no facing directions were verbally given to the participants.

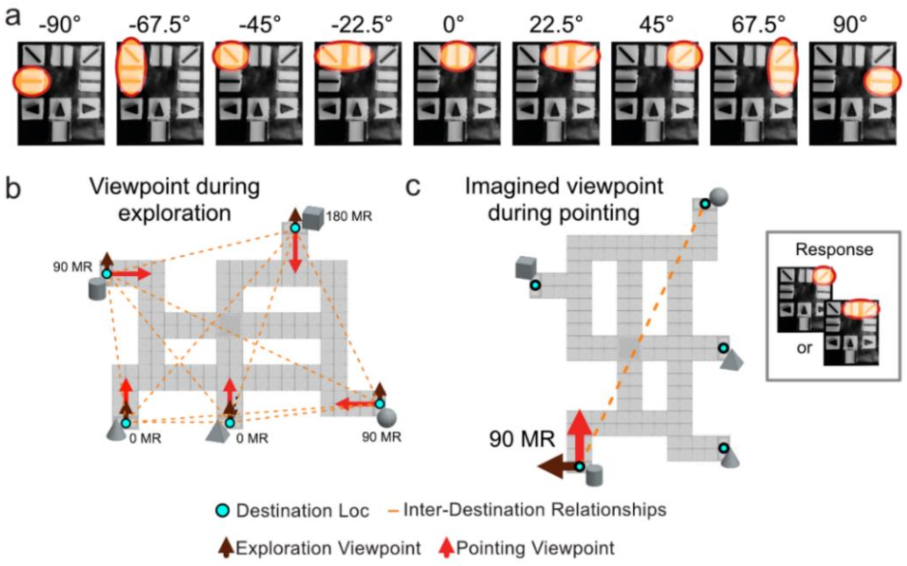


**Figure S6. Pointing task.** A. The keypad apparatus allowed 9 possible answers. B. The maze layout, including the locations of all destinations (sphere, cube, pyramid, cone and cylinder), the exploration viewpoints (as experienced when learning the tactile maze), the viewpoints from each destination (used in the JRD pointing task), and the spatial relationships (or directions and distances) between all pairs of destinations (e.g., directions to compare with participants' pointing data). C. The same maze layout viewed from the imagined viewpoint of an example trial (pointing from cylinder to sphere).

A direction estimate was provided on 5,369 of the 5,835 trials. Of these, 74 (1.4%) fell outside the ±90° range and were excluded, totaling 5,295 trials analyzed. Six participants (9.7%; 3 EB, 1 LB, 2 SC; 576 trials) used a joystick (before the apparatus was changed to the custom keypad). 30 of the 74 excluded trials came from joystick users. We tested the impact of the joystick on pointing scores (by exclusion and keypad-joystick difference, adjusted for group, age and sex; Table S2). The device did not predict pointing accuracy, and pointing-based effects held without joystick users. The reported analysis is therefore conservative by including joystick users.

##### Table S2. Device effect on pointing scores.

| Pointing (Group × block interaction, in manuscript, n=62) | Pointing (Group × block interaction, keypad only, n=56) | Device effect, β (keypad - joystick) |
|---|---|---|
| $\chi^2(2) = 3.954$, $p = .138$ | $\chi^2(2) = 1.47$, $p = .478$ | $\beta = -5.05$ (SE 3.27), $t(56) = -1.55$, $p = .128$ |

#### Text S4 – Pace analysis

The time required to reach a destination serves as an index of navigation efficiency. All trials are normalized according to their own straight-line distance (Euclidean distance between the start and goal locations), giving a measure of *pace* (in seconds per tile). However, the pace can only be computed on successful trials (arrival at destination; 3,054/5,835 trials, 52.3%), and this rate differs between groups and rises across blocks (see Fig. S4, left panel). Therefore, the time data is available only for a subset of trials. For the analysis, each trial was associated with a probability of arrival. Each timed trial then carried a weight equal to the inverse of that probability and stood in for the similar trials that produced no data. This correction assumes that arrival is unrelated to pace once block, group, route length, age and sex are held constant.

The probability of arrival came from a logistic model of arrival on block × group, route length, age and sex, fitted by generalized estimating equations. Weights were stabilized by the overall arrival rate and truncated at the 1st and 99th percentiles (0.59 and 3.18; totaling 61 weights modified). Balance is checked by comparing three distributions: all attempted trials, the trials that produced a time, and those same trials under their weights (see Table S3). Three quantities describe the result: the arrival model separates arrivals from non-arrivals with AUC = 0.700 (probability that a randomly chosen arrival receives a higher modelled probability than a randomly chosen non-arrival); the Kish effective sample size (number of equally weighted trials that would give the same precision) is 2,507 of 3,054 trials (82%); and the largest standardized difference between the analyzed trials and all attempted trials falls from 0.26 to 0.06. Pace is then fitted by weighted least squares with standard errors clustered by participant. Weighting moves a group mean pace by at most 0.028 s per tile. The LME models on this pace measure are presented in Tables S4-S9 or see the full analysis code in the dataset associated with the main manuscript.

**Table S3. Covariate balance.** "All trials" (average over every navigation trial, or the set the analysis is meant to represent), "Observed" (average over the timed 3,054 trials, counted once), "Weighted" (average over the 3,054 trials, each counted according to its weight). The last three rows are the proportion of trials contributed by each group. Standardized mean difference (SMD): gap between [observed or weighted] and all trials, divided by the standard deviation, $SD_{(\text{all trials})}$. The last two columns in the block row show that weighting takes a gap of $SMD_{(\text{observed})}$ = 0.261 to $SMD_{(\text{weighted})}$ = 0.015, and the largest remaining difference across all six variables is 0.06.

| *Variable* | *All trials (n=5,835)* | *Observed (n=3,054)* | *Weighted (n=3,054)* | *SD (all trials)* | *SMD (observed)* | *SMD (weighted)* |
|---|---|---|---|---|---|---|
| *Block* | 6.403 | 7.294 | 6.453 | 3.415 | +0.261 | +0.015 |
| *Route length (tiles)* | 16.914 | 16.565 | 16.815 | 3.514 | −0.099 | −0.028 |
| *Age* | 39.408 | 37.593 | 39.992 | 12.639 | −0.144 | +0.046 |
| *Proportion (EB)* | 0.259 | 0.231 | 0.286 | 0.438 | −0.065 | +0.062 |
| *Proportion (LB)* | 0.273 | 0.248 | 0.248 | 0.445 | −0.056 | −0.057 |
| *Proportion (SC)* | 0.468 | 0.521 | 0.466 | 0.499 | +0.107 | −0.004 |

#### Text S5 – LME models and statistical analyses

Six functions f(b) of the block index b = 1, …, 12, entered in place of a raw block term.

Linear: $f(b) = b - 1$
Logarithmic: $f(b) = \ln(b)$
Power: $f(b) = 1 - b^{(-1/2)}$
Exponential: $f(b) = 1 - e^{(-(b-1)/\tau)}$
Quadratic: $f(b) = (b - 1)$ and $(b - 1)^2$
Factor: $f(b) = \{1[b = k]\}, k = 2, \ldots, 12$

**Table S4 – Learning curve selection for the group comparisons (all groups modeled together).** Six candidate forms fitted for every dependent variable. ΔAICc is relative to the best form in that block. The time constant τ of the exponential form is not estimated inside the mixed model; it is profiled on the grid {2, 3, 4, 6, 8} and the best value carried into the comparison, reported in the τ column. Pace rows are fitted with weighted least squares and carry one parameter fewer than the mixed-model rows (no random-effect variance).

| *Outcome* | *Form* | *τ* | *n obs* | *k* | *logLik* | *AICc* | *ΔAICc* | *Conv.* |
|---|---|---|---|---|---|---|---|---|
| | | | ***Whole sample*** | | | | | |
| *Navigation performance* | exponential | 4.0 | 5835 | 11 | -29704.5 | 59431.0 | 0.0 | yes |
| | log | 4.0 | 5835 | 11 | -29706.8 | 59435.6 | 4.6 | yes |
| | power | 4.0 | 5835 | 11 | -29715.0 | 59452.1 | 21.1 | yes |
| | quadratic | 4.0 | 5835 | 12 | -29722.2 | 59468.5 | 37.4 | yes |
| | factor | 4.0 | 5835 | 39 | -29699.1 | 59476.8 | 45.7 | yes |
| | linear | 4.0 | 5835 | 11 | -29756.3 | 59534.6 | 103.6 | yes |
| *Pointing score* | power | 3.0 | 5295 | 11 | -22495.1 | 45012.3 | 0.0 | yes |
| | exponential | 3.0 | 5295 | 11 | -22497.6 | 45017.2 | 4.9 | yes |
| | log | 3.0 | 5295 | 11 | -22497.9 | 45017.8 | 5.5 | yes |
| | quadratic | 3.0 | 5295 | 12 | -22509.6 | 45043.3 | 30.9 | yes |
| | factor | 3.0 | 5295 | 39 | -22486.9 | 45052.4 | 40.1 | yes |
| | linear | 3.0 | 5295 | 11 | -22515.8 | 45053.7 | 41.4 | yes |
| *Pace (s per tile)* | exponential | 6.0 | 3054 | 9 | -890.8 | 1799.7 | 0.0 | yes |
| | log | 4.0 | 3054 | 9 | -891.2 | 1800.5 | 0.8 | yes |
| | quadratic | 4.0 | 3054 | 12 | -890.9 | 1805.8 | 6.1 | yes |
| | linear | 4.0 | 3054 | 9 | -896.0 | 1810.1 | 10.4 | yes |
| | power | 4.0 | 3054 | 9 | -896.6 | 1811.2 | 11.5 | yes |
| | factor | 4.0 | 3054 | 39 | -876.5 | 1832.0 | 32.3 | yes |
| | | | ***High performers*** | | | | | |
| *Navigation performance* | exponential | 4.0 | 4547 | 11 | -23053.9 | 46130.0 | 0.0 | yes |
| | log | 4.0 | 4547 | 11 | -23055.9 | 46133.8 | 3.9 | yes |
| | quadratic | 4.0 | 4547 | 12 | -23064.7 | 46153.4 | 23.5 | yes |
| | power | 4.0 | 4547 | 11 | -23065.8 | 46153.6 | 23.7 | yes |
| | factor | 4.0 | 4547 | 39 | -23047.9 | 46174.4 | 44.5 | yes |
| | linear | 4.0 | 4547 | 11 | -23100.1 | 46222.3 | 92.4 | yes |
| | linear | 3.0 | 4547 | 11 | -20669.2 | 41360.4 | 123.2 | yes |
| *Pointing score* | power | 2.0 | 4219 | 11 | -17566.3 | 35154.6 | 0.0 | yes |
| | exponential | 2.0 | 4219 | 11 | -17569.9 | 35161.9 | 7.3 | yes |
| | log | 2.0 | 4219 | 11 | -17570.1 | 35162.3 | 7.7 | yes |
| | quadratic | 2.0 | 4219 | 12 | -17582.5 | 35189.0 | 34.4 | yes |
| | factor | 2.0 | 4219 | 39 | -17564.3 | 35207.4 | 52.8 | yes |
| | linear | 2.0 | 4219 | 11 | -17690.9 | 35403.9 | 249.3 | yes |
| *Pace (s per tile)* | log | 4.0 | 2811 | 9 | -730.5 | 1479.1 | 0.0 | yes |
| | exponential | 6.0 | 2811 | 9 | -730.6 | 1479.2 | 0.1 | yes |
| | quadratic | 4.0 | 2811 | 12 | -729.4 | 1482.9 | 3.8 | yes |
| | power | 4.0 | 2811 | 9 | -735.8 | 1489.6 | 10.5 | yes |
| | linear | 4.0 | 2811 | 9 | -737.6 | 1493.3 | 14.2 | yes |
| | factor | 4.0 | 2811 | 39 | -716.4 | 1511.8 | 32.7 | yes |
| | | | ***Low performers*** | | | | | |
| *Navigation performance* | quadratic | 3.0 | 1288 | 12 | -6607.2 | 13238.6 | 0.0 | yes |
| | exponential | 3.0 | 1288 | 11 | -6609.7 | 13241.7 | 3.1 | yes |
| | power | 3.0 | 1288 | 11 | -6609.9 | 13242.1 | 3.5 | yes |

| | | | | | | | | |
|---|---|---|---|---|---|---|---|---|
| | linear | 3.0 | 1288 | 11 | -6613.5 | 13249.3 | 10.7 | yes |
| | factor | 3.0 | 1288 | 39 | -6589.9 | 13260.2 | 21.6 | yes |
| | log | 3.0 | 1288 | 11 | -6619.8 | 13261.8 | 23.2 | yes |
| *Pointing score* | power | 2.0 | 1076 | 11 | -4819.0 | 9660.2 | 0.0 | yes |
| | log | 2.0 | 1076 | 11 | -4819.1 | 9660.5 | 0.3 | yes |
| | exponential | 2.0 | 1076 | 11 | -4819.2 | 9660.6 | 0.4 | yes |
| | linear | 2.0 | 1076 | 11 | -4820.0 | 9662.3 | 2.1 | yes |
| | quadratic | 2.0 | 1076 | 12 | -4819.6 | 9663.5 | 3.2 | yes |
| | factor | 2.0 | 1076 | 39 | -4800.8 | 9682.6 | 22.4 | yes |
| *Pace (s per tile)* | exponential | 2.0 | 243 | 9 | -62.1 | 143.0 | 0.0 | yes |
| | power | 4.0 | 243 | 9 | -62.2 | 143.2 | 0.2 | yes |
| | log | 4.0 | 243 | 9 | -62.4 | 143.5 | 0.6 | yes |
| | linear | 4.0 | 243 | 9 | -62.8 | 144.4 | 1.5 | yes |
| | quadratic | 4.0 | 243 | 12 | -62.1 | 149.6 | 6.7 | yes |
| | factor | 4.0 | 243 | 38 | -48.3 | 187.2 | 44.3 | yes |

### Table S5 – Learning curve selection for each group separately (whole sample).

| *Outcome* | *Group* | *Form* | *τ* | *n obs* | *AICc* | *ΔAICc* | *Conv.* |
|---|---|---|---|---|---|---|---|
| *Navigation performance* | EB | quadratic | 3.0 | 1513 | 15506.5 | 0.0 | yes |
| | EB | exponential | 3.0 | 1513 | 15506.9 | 0.5 | yes |
| | EB | log | 3.0 | 1513 | 15509.0 | 2.5 | yes |
| | EB | power | 3.0 | 1513 | 15509.7 | 3.2 | yes |
| | EB | factor | 3.0 | 1513 | 15512.4 | 6.0 | yes |
| | EB | linear | 3.0 | 1513 | 15528.1 | 21.7 | yes |
| | LB | log | 3.0 | 1592 | 16340.3 | 0.0 | yes |
| | LB | power | 3.0 | 1592 | 16340.5 | 0.2 | yes |
| | LB | exponential | 3.0 | 1592 | 16344.4 | 4.1 | yes |
| | LB | factor | 3.0 | 1592 | 16354.1 | 13.8 | yes |
| | LB | quadratic | 3.0 | 1592 | 16357.9 | 17.6 | yes |
| | LB | linear | 3.0 | 1592 | 16362.8 | 22.5 | yes |
| | SC | exponential | 4.0 | 2730 | 27570.6 | 0.0 | yes |
| | SC | log | 4.0 | 2730 | 27578.3 | 7.7 | yes |
| | SC | quadratic | 4.0 | 2730 | 27593.8 | 23.2 | yes |
| | SC | power | 4.0 | 2730 | 27597.0 | 26.3 | yes |
| | SC | factor | 4.0 | 2730 | 27601.7 | 31.1 | yes |
| | SC | linear | 4.0 | 2730 | 27641.5 | 70.8 | yes |
| *Pointing score* | EB | quadratic | 2.0 | 1376 | 12152.8 | 0.0 | yes |
| | EB | factor | 2.0 | 1376 | 12153.5 | 0.6 | yes |
| | EB | power | 2.0 | 1376 | 12154.7 | 1.8 | yes |
| | EB | log | 2.0 | 1376 | 12155.2 | 2.4 | yes |
| | EB | exponential | 2.0 | 1376 | 12155.2 | 2.4 | yes |
| | EB | linear | 2.0 | 1376 | 12156.9 | 4.0 | yes |
| | LB | power | 2.0 | 1442 | 12313.1 | 0.0 | yes |
| | LB | log | 2.0 | 1442 | 12318.7 | 5.6 | yes |
| | LB | exponential | 2.0 | 1442 | 12320.5 | 7.5 | yes |
| | LB | quadratic | 2.0 | 1442 | 12334.2 | 21.2 | yes |
| | LB | linear | 2.0 | 1442 | 12336.2 | 23.1 | yes |
| | LB | factor | 2.0 | 1442 | 12336.3 | 23.3 | yes |
| | SC | exponential | 3.0 | 2477 | 20372.6 | 0.0 | yes |
| | SC | log | 3.0 | 2477 | 20375.0 | 2.4 | yes |
| | SC | power | 3.0 | 2477 | 20375.1 | 2.5 | yes |
| | SC | quadratic | 3.0 | 2477 | 20390.7 | 18.1 | yes |
| | SC | factor | 3.0 | 2477 | 20403.6 | 31.0 | yes |
| | SC | linear | 3.0 | 2477 | 20508.7 | 136.1 | yes |
| *Pace (s per tile)* | EB | exponential | 3.0 | 705 | 482.1 | 0.0 | yes |
| | EB | log | 4.0 | 705 | 482.5 | 0.4 | yes |
| | EB | power | 4.0 | 705 | 482.8 | 0.6 | yes |
| | EB | linear | 4.0 | 705 | 483.2 | 1.1 | yes |
| | EB | quadratic | 4.0 | 705 | 484.1 | 2.0 | yes |
| | EB | factor | 4.0 | 705 | 497.2 | 15.0 | yes |
| | LB | exponential | 8.0 | 757 | 413.5 | 0.0 | yes |
| | LB | linear | 4.0 | 757 | 413.6 | 0.1 | yes |
| | LB | quadratic | 4.0 | 757 | 415.1 | 1.6 | yes |
| | LB | log | 4.0 | 757 | 416.3 | 2.8 | yes |
| | LB | power | 4.0 | 757 | 420.8 | 7.3 | yes |
| | LB | factor | 4.0 | 757 | 421.6 | 8.1 | yes |
| | SC | log | 4.0 | 1592 | 827.9 | 0.0 | yes |
| | SC | exponential | 4.0 | 1592 | 829.5 | 1.6 | yes |
| | SC | quadratic | 4.0 | 1592 | 833.5 | 5.6 | yes |
| | SC | power | 4.0 | 1592 | 835.2 | 7.3 | yes |
| | SC | factor | 4.0 | 1592 | 839.6 | 11.7 | yes |
| | SC | linear | 4.0 | 1592 | 840.8 | 12.9 | yes |

### Table S6 – Learning curve selection for each group separately (high performers)

| *Outcome* | *Group* | *Form* | *τ* | *n obs* | *AICc* | *ΔAICc* | *Conv.* |
|---|---|---|---|---|---|---|---|
| *Navigation performance* | EB | quadratic | 4.0 | 953 | 9728.3 | 0.0 | yes |
| | EB | exponential | 4.0 | 953 | 9728.9 | 0.6 | yes |
| | EB | power | 4.0 | 953 | 9732.5 | 4.2 | yes |
| | EB | factor | 4.0 | 953 | 9741.4 | 13.1 | yes |
| | EB | linear | 4.0 | 953 | 9743.8 | 15.5 | yes |
| | EB | log | 4.0 | 953 | 9744.7 | 16.4 | yes |
| | LB | log | 4.0 | 1248 | 12784.9 | 0.0 | yes |
| | LB | power | 4.0 | 1248 | 12787.0 | 2.1 | yes |
| | LB | exponential | 4.0 | 1248 | 12789.4 | 4.5 | yes |
| | LB | quadratic | 4.0 | 1248 | 12791.4 | 6.5 | yes |
| | LB | factor | 4.0 | 1248 | 12792.1 | 7.2 | yes |
| | LB | linear | 4.0 | 1248 | 12799.5 | 14.6 | yes |
| | SC | exponential | 4.0 | 2346 | 23605.5 | 0.0 | yes |
| | SC | log | 4.0 | 2346 | 23613.8 | 8.3 | yes |
| | SC | quadratic | 4.0 | 2346 | 23626.9 | 21.4 | yes |
| | SC | power | 4.0 | 2346 | 23629.6 | 24.0 | yes |
| | SC | factor | 4.0 | 2346 | 23635.6 | 30.1 | yes |
| | SC | linear | 4.0 | 2346 | 23678.7 | 73.2 | yes |
| *Pointing score* | EB | quadratic | 2.0 | 897 | 7773.8 | 0.0 | yes |
| | EB | power | 2.0 | 897 | 7775.4 | 1.6 | yes |
| | EB | exponential | 2.0 | 897 | 7776.0 | 2.2 | yes |
| | EB | log | 2.0 | 897 | 7776.0 | 2.3 | yes |
| | EB | linear | 2.0 | 897 | 7777.6 | 3.8 | yes |
| | EB | factor | 2.0 | 897 | 7778.8 | 5.0 | yes |
| | LB | power | 2.0 | 1162 | 9705.2 | 0.0 | yes |
| | LB | log | 2.0 | 1162 | 9711.2 | 6.0 | yes |
| | LB | exponential | 2.0 | 1162 | 9712.8 | 7.6 | yes |
| | LB | quadratic | 2.0 | 1162 | 9728.8 | 23.5 | yes |
| | LB | linear | 2.0 | 1162 | 9731.3 | 26.0 | yes |
| | LB | factor | 2.0 | 1162 | 9735.3 | 30.0 | yes |
| | SC | exponential | 3.0 | 2160 | 17590.3 | 0.0 | yes |
| | SC | power | 3.0 | 2160 | 17591.3 | 1.0 | yes |
| | SC | log | 3.0 | 2160 | 17592.5 | 2.2 | yes |
| | SC | quadratic | 3.0 | 2160 | 17602.5 | 12.2 | yes |
| | SC | factor | 3.0 | 2160 | 17614.2 | 23.9 | yes |
| | SC | linear | 3.0 | 2160 | 17710.7 | 120.4 | yes |
| *Pace (s per tile)* | EB | exponential | 2.0 | 593 | 341.1 | 0.0 | yes |
| | EB | power | 4.0 | 593 | 342.1 | 1.0 | yes |
| | EB | quadratic | 4.0 | 593 | 342.4 | 1.3 | yes |
| | EB | log | 4.0 | 593 | 342.5 | 1.4 | yes |
| | EB | linear | 4.0 | 593 | 345.5 | 4.4 | yes |
| | EB | factor | 4.0 | 593 | 356.7 | 15.6 | yes |
| | LB | exponential | 8.0 | 696 | 315.5 | 0.0 | yes |
| | LB | linear | 4.0 | 696 | 316.9 | 1.5 | yes |
| | LB | quadratic | 4.0 | 696 | 317.0 | 1.5 | yes |
| | LB | log | 4.0 | 696 | 317.1 | 1.6 | yes |
| | LB | power | 4.0 | 696 | 320.5 | 5.0 | yes |
| | LB | factor | 4.0 | 696 | 327.8 | 12.3 | yes |
| | SC | log | 4.0 | 1522 | 758.1 | 0.0 | yes |
| | SC | exponential | 6.0 | 1522 | 759.4 | 1.2 | yes |
| | SC | quadratic | 4.0 | 1522 | 763.3 | 5.2 | yes |
| | SC | power | 4.0 | 1522 | 767.3 | 9.2 | yes |
| | SC | factor | 4.0 | 1522 | 767.4 | 9.3 | yes |
| | SC | linear | 4.0 | 1522 | 770.1 | 12.0 | yes |

#### Table S7 – Learning curve selection for each group separately (low performers)

| *Outcome* | *Group* | *Form* | *τ* | *n obs* | *AICc* | *ΔAICc* | *Conv.* |
|---|---|---|---|---|---|---|---|
| *Navigation performance* | EB | quadratic | 2.0 | 560 | 5759.4 | 0.0 | yes |
| | EB | power | 2.0 | 560 | 5764.5 | 5.1 | yes |
| | EB | exponential | 2.0 | 560 | 5765.7 | 6.3 | yes |
| | EB | log | 2.0 | 560 | 5766.2 | 6.9 | yes |
| | EB | factor | 2.0 | 560 | 5767.3 | 7.9 | yes |
| | EB | linear | 2.0 | 560 | 5771.2 | 11.8 | yes |
| | LB | quadratic | 2.0 | 344 | 3540.1 | 0.0 | yes |
| | LB | power | 2.0 | 344 | 3541.7 | 1.6 | yes |
| | LB | log | 2.0 | 344 | 3542.6 | 2.5 | yes |
| | LB | factor | 2.0 | 344 | 3543.0 | 2.9 | yes |
| | LB | exponential | 2.0 | 344 | 3543.7 | 3.6 | yes |
| | LB | linear | 2.0 | 344 | 3544.4 | 4.3 | yes |
| | SC | quadratic | 8.0 | 384 | 3936.0 | 0.0 | yes |
| | SC | log | 8.0 | 384 | 3938.8 | 2.9 | yes |
| | SC | exponential | 8.0 | 384 | 3938.8 | 2.9 | yes |
| | SC | linear | 8.0 | 384 | 3939.1 | 3.1 | yes |
| | SC | power | 8.0 | 384 | 3945.4 | 9.5 | yes |
| | SC | factor | 8.0 | 384 | 3947.2 | 11.2 | yes |
| *Pointing score* | EB | quadratic | 3.0 | 479 | 4358.3 | 0.0 | yes |
| | EB | exponential | 3.0 | 479 | 4359.9 | 1.6 | yes |
| | EB | log | 3.0 | 479 | 4360.0 | 1.8 | yes |
| | EB | power | 3.0 | 479 | 4360.2 | 1.9 | yes |
| | EB | linear | 3.0 | 479 | 4360.5 | 2.2 | yes |
| | EB | factor | 3.0 | 479 | 4368.6 | 10.3 | yes |
| | LB | quadratic | 2.0 | 280 | 2536.9 | 0.0 | yes |
| | LB | factor | 2.0 | 280 | 2537.7 | 0.8 | yes |
| | LB | log | 2.0 | 280 | 2538.5 | 1.6 | yes |
| | LB | exponential | 2.0 | 280 | 2538.9 | 2.0 | yes |
| | LB | linear | 2.0 | 280 | 2539.2 | 2.3 | yes |
| | LB | power | 2.0 | 280 | 2539.3 | 2.4 | yes |
| | SC | exponential | 8.0 | 317 | 2744.8 | 0.0 | yes |
| | SC | linear | 8.0 | 317 | 2745.4 | 0.5 | yes |
| | SC | power | 8.0 | 317 | 2746.7 | 1.9 | yes |
| | SC | log | 8.0 | 317 | 2747.7 | 2.9 | yes |
| | SC | quadratic | 8.0 | 317 | 2751.8 | 7.0 | yes |
| | SC | factor | 8.0 | 317 | 2760.6 | 15.8 | yes |
| *Pace (s per tile)* | EB | linear | 4.0 | 112 | 58.7 | 0.0 | yes |
| | EB | exponential | 8.0 | 112 | 58.7 | 0.1 | yes |
| | EB | log | 4.0 | 112 | 58.8 | 0.2 | yes |
| | EB | power | 4.0 | 112 | 58.9 | 0.2 | yes |
| | EB | quadratic | 4.0 | 112 | 60.7 | 2.0 | yes |
| | EB | factor | 4.0 | 112 | 72.1 | 13.4 | yes |
| | LB | exponential | 2.0 | 61 | 42.4 | 0.0 | yes |
| | LB | power | 4.0 | 61 | 42.4 | 0.1 | yes |
| | LB | log | 4.0 | 61 | 43.3 | 1.0 | yes |
| | LB | linear | 4.0 | 61 | 45.0 | 2.6 | yes |
| | LB | quadratic | 4.0 | 61 | 46.4 | 4.0 | yes |
| | LB | factor | 4.0 | 61 | 66.0 | 23.6 | yes |
| | SC | exponential | 3.0 | 70 | 35.3 | 0.0 | yes |
| | SC | power | 4.0 | 70 | 35.5 | 0.2 | yes |
| | SC | log | 4.0 | 70 | 35.7 | 0.4 | yes |
| | SC | linear | 4.0 | 70 | 36.5 | 1.2 | yes |
| | SC | quadratic | 4.0 | 70 | 37.6 | 2.4 | yes |
| | SC | factor | 4.0 | 70 | 51.6 | 16.3 | yes |

**Table S8 – Omnibus tests, for all dependent variables and performance classifications.** Every omnibus test behind the analyses reported in the manuscript, together with the outcome-definition checks of Table S-A. The form column gives the block term selected by AICc for that dependent variable and stratum. The pace rows come from the weighted analysis: all three tests are joint Wald tests on the participant-clustered covariance of the weighted least-squares fit, since that estimator has no likelihood to compare.

| ***Outcome*** | ***Stratum*** | ***Form*** | ***Omnibus test*** | ***Test*** | $\chi^2$ | ***df*** | ***p*** |
|---|---|---|---|---|---|---|---|
| *Navigation performance* | whole sample | exponential | Group x Block | LRT | 8.77 | 2 | .012 * |
| | whole sample | exponential | Group | LRT | 1.13 | 2 | .568 |
| | whole sample | exponential | Group at baseline | Wald | 0.25 | 2 | .881 |
| | high | exponential | Group x Block | LRT | 3.28 | 2 | .194 |
| | high | exponential | Group | LRT | 1.48 | 2 | .477 |
| | high | exponential | Group at baseline | Wald | 1.64 | 2 | .441 |
| | low | quadratic | Group x Block | LRT | 11.06 | 4 | .026 * |
| | low | quadratic | Group | LRT | 0.17 | 2 | .920 |
| | low | quadratic | Group at baseline | Wald | 0.97 | 2 | .615 |
| *Pointing score* | whole sample | power | Group x Block | LRT | 3.95 | 2 | .138 |
| | whole sample | power | Group | LRT | 10.57 | 2 | .005 * |
| | whole sample | power | Group at baseline | Wald | 3.56 | 2 | .169 |
| | high | power | Group x Block | LRT | 4.13 | 2 | .127 |
| | high | power | Group | LRT | 5.96 | 2 | .051 |
| | high | power | Group at baseline | Wald | 2.64 | 2 | .267 |
| | low | power | Group x Block | LRT | 0.24 | 2 | .886 |
| | low | power | Group | LRT | 1.32 | 2 | .516 |
| | low | power | Group at baseline | Wald | 0.31 | 2 | .857 |
| *Pace (s per tile)* | whole sample | exponential | Group x Block | Wald | 14.33 | 2 | < .001 * |
| | whole sample | exponential | Group | Wald | 0.76 | 2 | .685 |
| | whole sample | exponential | Group at baseline | Wald | 7.08 | 2 | .029 * |
| | high | log | Group x Block | Wald | 13.90 | 2 | .001 * |
| | high | log | Group | Wald | 1.54 | 2 | .462 |
| | high | log | Group at baseline | Wald | 9.79 | 2 | .007 * |
| | low | exponential | Group x Block | Wald | 7.22 | 2 | .027 * |
| | low | exponential | Group | Wald | 3.37 | 2 | .185 |
| | low | exponential | Group at baseline | Wald | 6.09 | 2 | .048 * |

**Table S9 – Contrasts and within-group slopes, all dependent variables and performance classifications.** Coefficients, standard errors, z, 95% confidence intervals and Holm-corrected p for every contrast and within-group slope. Baseline contrasts are group differences at block 1. Where the quadratic form was selected the block term contributes two coefficients, so a group has a linear and a curvature coefficient rather than a single slope, and these are corrected within their own family. The pace rows come from the weighted least-squares fit with the inverse-probability weights; their confidence intervals are given to two decimals because pace is measured in seconds per tile.

| *Outcome* | *Stratum* | *Effect* | *Contrast* | *β* | *SE* | *z* | *95% CI* | *p* | *p Holm* |
|---|---|---|---|---|---|---|---|---|---|
| *Navigation performance* | whole sample | overall group difference | LB - EB | -1.34 | 6.96 | -0.19 | [-15.0, 12.3] | .847 | .954 |
| | whole sample | overall group difference | SC - EB | 5.23 | 6.66 | 0.78 | [-7.8, 18.3] | .432 | .954 |
| | whole sample | overall group difference | SC - LB | 6.57 | 6.57 | 1.00 | [-6.3, 19.5] | .318 | .954 |
| | whole sample | baseline (block 1) difference | LB - EB | -3.72 | 7.41 | -0.50 | [-18.2, 10.8] | .615 | 1.000 |
| | whole sample | baseline (block 1) difference | SC - EB | -1.72 | 6.68 | -0.26 | [-14.8, 11.4] | .797 | 1.000 |
| | whole sample | baseline (block 1) difference | SC - LB | 2.01 | 6.80 | 0.30 | [-11.3, 15.3] | .768 | 1.000 |
| | whole sample | learning-rate difference (slope) | LB - EB | 6.59 | 7.36 | 0.90 | [-7.8, 21.0] | .371 | .371 |
| | whole sample | learning-rate difference (slope) | SC - EB | 19.18 | 6.57 | 2.92 | [6.3, 32.1] | .004 | .011 * |
| | whole sample | learning-rate difference (slope) | SC - LB | 12.59 | 6.46 | 1.95 | [-0.1, 25.3] | .051 | .103 |
| | whole sample | within-group slope | EB | 34.72 | 5.27 | 6.59 | [24.4, 45.1] | < .001 | < .001 * |
| | whole sample | within-group slope | LB | 41.31 | 5.13 | 8.05 | [31.3, 51.4] | < .001 | < .001 * |
| | whole sample | within-group slope | SC | 53.90 | 3.92 | 13.74 | [46.2, 61.6] | < .001 | < .001 * |
| | high | overall group difference | LB - EB | -5.88 | 6.15 | -0.96 | [-17.9, 6.2] | .339 | .777 |
| | high | overall group difference | SC - EB | 0.35 | 6.02 | 0.06 | [-11.4, 12.1] | .953 | .953 |
| | high | overall group difference | SC - LB | 6.24 | 5.53 | 1.13 | [-4.6, 17.1] | .259 | .777 |
| | high | baseline (block 1) difference | LB - EB | -9.96 | 8.78 | -1.13 | [-27.2, 7.3] | .257 | .738 |
| | high | baseline (block 1) difference | SC - EB | -9.43 | 8.13 | -1.16 | [-25.4, 6.5] | .246 | .738 |
| | high | baseline (block 1) difference | SC - LB | 0.53 | 7.51 | 0.07 | [-14.2, 15.2] | .943 | .943 |
| | high | learning-rate difference (slope) | LB - EB | 5.49 | 8.46 | 0.65 | [-11.1, 22.1] | .516 | .530 |
| | high | learning-rate difference (slope) | SC - EB | 13.18 | 7.54 | 1.75 | [-1.6, 28.0] | .081 | .242 |
| | high | learning-rate difference (slope) | SC - LB | 7.68 | 6.89 | 1.12 | [-5.8, 21.2] | .265 | .530 |
| | high | within-group slope | EB | 43.09 | 6.37 | 6.77 | [30.6, 55.6] | < .001 | < .001 * |
| | high | within-group slope | LB | 48.59 | 5.57 | 8.72 | [37.7, 59.5] | < .001 | < .001 * |
| | high | within-group slope | SC | 56.27 | 4.05 | 13.91 | [48.3, 64.2] | < .001 | < .001 * |
| | low | overall group difference | LB - EB | -2.62 | 7.07 | -0.37 | [-16.5, 11.2] | .711 | 1.000 |
| | low | overall group difference | SC - EB | -0.46 | 6.78 | -0.07 | [-13.7, 12.8] | .946 | 1.000 |

| | | | | | | | | | |
|---|---|---|---|---|---|---|---|---|---|
| | low | overall group difference | SC - LB | 2.16 | 6.46 | 0.33 | [-10.5, 14.8] | .738 | 1.000 |
| | low | baseline (block 1) difference | LB - EB | 2.67 | 9.45 | 0.28 | [-15.8, 21.2] | .777 | 1.000 |
| *Navigation performance* | low | baseline (block 1) difference | SC - EB | -6.33 | 9.23 | -0.69 | [-24.4, 11.8] | .493 | 1.000 |
| | low | baseline (block 1) difference | SC - LB | -9.00 | 9.43 | -0.95 | [-27.5, 9.5] | .340 | 1.000 |
| | low | learning-rate difference (linear) | LB - EB | -2.45 | 3.04 | -0.81 | [-8.4, 3.5] | .419 | 1.000 |
| | low | learning-rate difference (linear) | SC - EB | -1.17 | 2.93 | -0.40 | [-6.9, 4.6] | .690 | 1.000 |
| | low | learning-rate difference (linear) | SC - LB | 1.29 | 3.27 | 0.39 | [-5.1, 7.7] | .694 | 1.000 |
| | low | learning-rate difference (curvature) | LB - EB | 0.18 | 0.28 | 0.65 | [-0.4, 0.7] | .519 | 1.000 |
| | low | learning-rate difference (curvature) | SC - EB | 0.30 | 0.26 | 1.14 | [-0.2, 0.8] | .254 | .763 |
| | low | learning-rate difference (curvature) | SC - LB | 0.11 | 0.30 | 0.38 | [-0.5, 0.7] | .701 | 1.000 |
| | low | within-group curvature | EB | -0.53 | 0.17 | -3.14 | [-0.9, -0.2] | .002 | .005 * |
| | low | within-group curvature | LB | -0.35 | 0.22 | -1.57 | [-0.8, 0.1] | .117 | .235 |
| | low | within-group curvature | SC | -0.24 | 0.20 | -1.20 | [-0.6, 0.1] | .230 | .235 |
| | low | within-group linear | EB | 7.00 | 1.88 | 3.72 | [3.3, 10.7] | < .001 | < .001 * |
| | low | within-group linear | LB | 4.55 | 2.38 | 1.91 | [-0.1, 9.2] | .056 | .056 |
| | low | within-group linear | SC | 5.84 | 2.24 | 2.61 | [1.4, 10.2] | .009 | .018 * |
| *Pointing score* | whole sample | overall group difference | LB - EB | 3.12 | 2.39 | 1.31 | [-1.6, 7.8] | .191 | .191 |
| | whole sample | overall group difference | SC - EB | 7.14 | 2.14 | 3.33 | [2.9, 11.3] | < .001 | .003 * |
| | whole sample | overall group difference | SC - LB | 4.02 | 2.15 | 1.87 | [-0.2, 8.2] | .061 | .122 |
| | whole sample | baseline (block 1) difference | LB - EB | -0.86 | 3.13 | -0.27 | [-7.0, 5.3] | .784 | .784 |
| | whole sample | baseline (block 1) difference | SC - EB | 3.93 | 2.81 | 1.40 | [-1.6, 9.4] | .162 | .325 |
| | whole sample | baseline (block 1) difference | SC - LB | 4.78 | 2.81 | 1.70 | [-0.7, 10.3] | .088 | .265 |
| | whole sample | learning-rate difference (slope) | LB - EB | 8.94 | 4.73 | 1.89 | [-0.3, 18.2] | .059 | .176 |
| | whole sample | learning-rate difference (slope) | SC - EB | 7.21 | 4.22 | 1.71 | [-1.1, 15.5] | .087 | .176 |
| | whole sample | learning-rate difference (slope) | SC - LB | -1.73 | 4.12 | -0.42 | [-9.8, 6.3] | .675 | .675 |
| | whole sample | within-group slope | EB | 7.29 | 3.41 | 2.14 | [0.6, 14.0] | .032 | .032 * |
| | whole sample | within-group slope | LB | 16.23 | 3.28 | 4.95 | [9.8, 22.7] | < .001 | < .001 * |
| | whole sample | within-group slope | SC | 14.50 | 2.49 | 5.83 | [9.6, 19.4] | < .001 | < .001 * |
| | high | overall group difference | LB - EB | 2.15 | 2.32 | 0.93 | [-2.4, 6.7] | .354 | .354 |
| | high | overall group difference | SC - EB | 5.31 | 2.19 | 2.43 | [1.0, 9.6] | .015 | .046 * |
| | high | overall group difference | SC - LB | 3.16 | 2.00 | 1.58 | [-0.8, 7.1] | .113 | .227 |
| | high | baseline (block 1) difference | LB - EB | -3.07 | 3.39 | -0.90 | [-9.7, 3.6] | .366 | .732 |
| | high | baseline (block 1) difference | SC - EB | 1.61 | 3.12 | 0.52 | [-4.5, 7.7] | .606 | .732 |

| | | | | | | | | | |
|---|---|---|---|---|---|---|---|---|---|
| | high | baseline (block 1) difference | SC - LB | 4.67 | 2.88 | 1.62 | [-1.0, 10.3] | .105 | .314 |
| *Pointing score* | high | learning-rate difference (slope) | LB - EB | 11.37 | 5.58 | 2.04 | [0.4, 22.3] | .042 | .125 |
| | high | learning-rate difference (slope) | SC - EB | 8.08 | 4.97 | 1.63 | [-1.7, 17.8] | .104 | .208 |
| | high | learning-rate difference (slope) | SC - LB | -3.29 | 4.54 | -0.73 | [-12.2, 5.6] | .468 | .468 |
| | high | within-group slope | EB | 6.79 | 4.19 | 1.62 | [-1.4, 15.0] | .105 | .105 |
| | high | within-group slope | LB | 18.16 | 3.68 | 4.94 | [11.0, 25.4] | < .001 | < .001 * |
| | high | within-group slope | SC | 14.87 | 2.66 | 5.59 | [9.7, 20.1] | < .001 | < .001 * |
| | low | overall group difference | LB - EB | 1.53 | 4.73 | 0.32 | [-7.7, 10.8] | .746 | .807 |
| | low | overall group difference | SC - EB | 5.10 | 4.60 | 1.11 | [-3.9, 14.1] | .267 | .800 |
| | low | overall group difference | SC - LB | 3.57 | 4.27 | 0.84 | [-4.8, 11.9] | .403 | .807 |
| | low | baseline (block 1) difference | LB - EB | 0.77 | 6.28 | 0.12 | [-11.5, 13.1] | .902 | 1.000 |
| | low | baseline (block 1) difference | SC - EB | 3.15 | 6.00 | 0.52 | [-8.6, 14.9] | .600 | 1.000 |
| | low | baseline (block 1) difference | SC - LB | 2.37 | 5.97 | 0.40 | [-9.3, 14.1] | .691 | 1.000 |
| | low | learning-rate difference (slope) | LB - EB | 1.57 | 9.04 | 0.17 | [-16.1, 19.3] | .862 | 1.000 |
| | low | learning-rate difference (slope) | SC - EB | 4.28 | 8.62 | 0.50 | [-12.6, 21.2] | .620 | 1.000 |
| | low | learning-rate difference (slope) | SC - LB | 2.71 | 9.45 | 0.29 | [-15.8, 21.2] | .774 | 1.000 |
| | low | within-group slope | EB | 8.23 | 5.68 | 1.45 | [-2.9, 19.4] | .147 | .294 |
| | low | within-group slope | LB | 9.80 | 6.99 | 1.40 | [-3.9, 23.5] | .161 | .294 |
| | low | within-group slope | SC | 12.51 | 6.46 | 1.94 | [-0.1, 25.2] | .053 | .158 |
| *Pace (s per tile)* | whole sample | overall group difference | LB - EB | 0.03 | 0.07 | 0.37 | [-0.12, 0.17] | .711 | 1.000 |
| | whole sample | overall group difference | SC - EB | 0.05 | 0.06 | 0.81 | [-0.07, 0.18] | .417 | 1.000 |
| | whole sample | overall group difference | SC - LB | 0.02 | 0.05 | 0.48 | [-0.08, 0.13] | .633 | 1.000 |
| | whole sample | baseline (block 1) difference | LB - EB | 0.10 | 0.08 | 1.22 | [-0.06, 0.26] | .223 | .223 |
| | whole sample | baseline (block 1) difference | SC - EB | 0.20 | 0.08 | 2.52 | [0.04, 0.35] | .012 | .035 * |
| | whole sample | baseline (block 1) difference | SC - LB | 0.10 | 0.06 | 1.63 | [-0.02, 0.22] | .103 | .207 |
| | whole sample | learning-rate difference (slope) | LB - EB | -0.14 | 0.08 | -1.69 | [-0.30, 0.02] | .091 | .128 |
| | whole sample | learning-rate difference (slope) | SC - EB | -0.28 | 0.07 | -3.75 | [-0.42, -0.13] | < .001 | < .001 * |
| | whole sample | learning-rate difference (slope) | SC - LB | -0.14 | 0.07 | -1.85 | [-0.28, 0.01] | .064 | .128 |
| | whole sample | within-group slope | EB | -0.09 | 0.06 | -1.61 | [-0.21, 0.02] | .108 | .108 |
| | whole sample | within-group slope | LB | -0.23 | 0.06 | -4.03 | [-0.35, -0.12] | < .001 | < .001 * |
| | whole sample | within-group slope | SC | -0.37 | 0.04 | -8.41 | [-0.46, -0.28] | < .001 | < .001 * |
| | high | overall group difference | LB - EB | 0.04 | 0.07 | 0.60 | [-0.10, 0.18] | .546 | 1.000 |
| | high | overall group difference | SC - EB | 0.08 | 0.06 | 1.20 | [-0.05, 0.20] | .230 | .689 |

| | | | | | | | | | |
|---|---|---|---|---|---|---|---|---|---|
| | high | overall group difference | SC - LB | 0.03 | 0.05 | 0.64 | [-0.07, 0.14] | .520 | 1.000 |
| *Pace (s per tile)* | high | baseline (block 1) difference | LB - EB | 0.10 | 0.08 | 1.16 | [-0.07, 0.26] | .247 | .247 |
| | high | baseline (block 1) difference | SC - EB | 0.23 | 0.08 | 2.78 | [0.07, 0.40] | .005 | .016 * |
| | high | baseline (block 1) difference | SC - LB | 0.14 | 0.06 | 2.33 | [0.02, 0.25] | .020 | .040 * |
| | high | learning-rate difference (slope) | LB - EB | -0.03 | 0.03 | -1.09 | [-0.09, 0.03] | .276 | .276 |
| | high | learning-rate difference (slope) | SC - EB | -0.09 | 0.03 | -3.54 | [-0.15, -0.04] | < .001 | .001 * |
| | high | learning-rate difference (slope) | SC - LB | -0.06 | 0.03 | -2.26 | [-0.11, -0.01] | .024 | .048 * |
| | high | within-group slope | EB | -0.05 | 0.02 | -2.26 | [-0.09, -0.01] | .024 | .024 * |
| | high | within-group slope | LB | -0.08 | 0.02 | -3.73 | [-0.13, -0.04] | < .001 | < .001 * |
| | high | within-group slope | SC | -0.14 | 0.01 | -9.56 | [-0.17, -0.11] | < .001 | < .001 * |
| | low | overall group difference | LB - EB | -0.05 | 0.07 | -0.71 | [-0.19, 0.09] | .479 | .712 |
| | low | overall group difference | SC - EB | -0.15 | 0.09 | -1.72 | [-0.33, 0.02] | .085 | .255 |
| | low | overall group difference | SC - LB | -0.10 | 0.11 | -0.92 | [-0.32, 0.12] | .356 | .712 |
| | low | baseline (block 1) difference | LB - EB | -0.16 | 0.08 | -1.90 | [-0.33, 0.01] | .057 | .115 |
| | low | baseline (block 1) difference | SC - EB | 0.16 | 0.12 | 1.28 | [-0.08, 0.40] | .199 | .199 |
| | low | baseline (block 1) difference | SC - LB | 0.32 | 0.14 | 2.30 | [0.05, 0.59] | .022 | .065 |
| | low | learning-rate difference (slope) | LB - EB | 0.15 | 0.10 | 1.50 | [-0.04, 0.34] | .134 | .134 |
| | low | learning-rate difference (slope) | SC - EB | -0.35 | 0.18 | -1.92 | [-0.70, 0.01] | .055 | .111 |
| | low | learning-rate difference (slope) | SC - LB | -0.49 | 0.19 | -2.61 | [-0.86, -0.12] | .009 | .027 * |
| | low | within-group slope | EB | -0.02 | 0.06 | -0.32 | [-0.13, 0.09] | .749 | .749 |
| | low | within-group slope | LB | 0.13 | 0.08 | 1.56 | [-0.03, 0.29] | .120 | .239 |
| | low | within-group slope | SC | -0.36 | 0.17 | -2.12 | [-0.70, -0.03] | .034 | .102 |

#### Text S6 – Mental rotation analysis

Participants learned the tactile maze facing North: trials with a start location facing North (Pyramid and Cone) constitute the 0° MR level; trials facing East (cylinder) or West (sphere), the 90° level; and trials facing South (cube), the 180° level. The model was a mixed ANCOVA (62 participants x 3 MR levels) according to the following equation: PointingScore ~ Group × MR + Age + Sex, with a random intercept for participant, fitted by maximum likelihood.

##### Table S10. Pointing score by group and rotation level.

| *Group* | *Rotation* | *Participants* | *N trials* | *Mean (%)* | *SD* |
|---|---|---|---|---|---|
| *EB* | 0° | 16 | 505 | 83.75 | 5.38 |
| *EB* | 90° | 16 | 659 | 73.47 | 9.58 |
| *EB* | 180° | 16 | 212 | 70.32 | 14.33 |
| *LB* | 0° | 17 | 536 | 85.11 | 5.18 |
| *LB* | 90° | 17 | 689 | 77.22 | 10.18 |
| *LB* | 180° | 17 | 217 | 77.28 | 10.00 |
| *SC* | 0° | 29 | 916 | 86.17 | 7.44 |
| *SC* | 90° | 29 | 1201 | 83.80 | 8.15 |
| *SC* | 180° | 29 | 360 | 84.48 | 11.36 |

##### Table S11. Coefficients of the mixed ANCOVA, with EB and 0° as the reference levels.

| *Term* | *β* | *SE* | *z* | *95% CI* | *p* |
|---|---|---|---|---|---|
| *Intercept* | +87.240 | 4.319 | +20.200 | [78.78, 95.70] | < .001 |
| *Group: LB* | +1.798 | 3.131 | +0.574 | [−4.34, 7.93] | .566 |
| *Group: SC* | +1.897 | 2.820 | +0.673 | [−3.63, 7.42] | .501 |
| *MR: 90°* | −10.278 | 2.205 | −4.662 | [−14.60, −5.96] | < .001 |
| *MR: 180°* | −13.425 | 2.205 | −6.089 | [−17.75, −9.10] | < .001 |
| *Sex* | +2.294 | 1.938 | +1.184 | [−1.50, 6.09] | .237 |
| *LB × 90°* | +2.383 | 3.072 | +0.776 | [−3.64, 8.40] | .438 |
| *SC × 90°* | +7.909 | 2.747 | +2.880 | [2.53, 13.29] | .004 |
| *LB × 180°* | +5.592 | 3.072 | +1.820 | [−0.43, 11.61] | .069 |
| *SC × 180°* | +11.729 | 2.747 | +4.271 | [6.35, 17.11] | < .001 |
| *Age* | −0.117 | 0.080 | −1.463 | [−0.27, 0.04] | .144 |

##### Table S12. Omnibus tests.

| *Effect* | *Test* | $\chi^2$ | *df* | *p* |
|---|---|---|---|---|
| *Group x MR (interaction)* | LRT | 18.735 | 4 | < .001 |
| *Group (main effect)* | LRT | 11.936 | 2 | .003 |
| *MR (main effect)* | LRT | 30.749 | 2 | < .001 |
| *Group at MR = 0 (reference level)* | Wald | 0.513 | 2 | .774 |

Table S13. All contrasts from the model.

| *Family* | *Comparison* | *n* | *β* | *SE* | *z* | *95% CI* | *p* | *p Holm* |
|---|---|---|---|---|---|---|---|---|
| *Group at 0°* | EB vs LB | 33 | −1.798 | 3.131 | −0.574 | [−7.94, 4.34] | .566 | 1.000 |
| *Group at 0°* | EB vs SC | 45 | −1.897 | 2.820 | −0.673 | [−7.42, 3.63] | .501 | 1.000 |
| *Group at 0°* | LB vs SC | 46 | −0.099 | 2.848 | −0.035 | [−5.68, 5.48] | .972 | 1.000 |
| *Group at 180°* | EB vs LB | 33 | −7.389 | 3.131 | −2.360 | [−13.53, −1.25] | .018 | .037 |
| *Group at 180°* | EB vs SC | 45 | −13.626 | 2.820 | −4.832 | [−19.15, −8.10] | < .001 | < .001 |
| *Group at 180°* | LB vs SC | 46 | −6.237 | 2.848 | −2.190 | [−11.82, −0.66] | .028 | .037 |
| *Group at 90°* | EB vs LB | 33 | −4.181 | 3.131 | −1.335 | [−10.32, 1.96] | .182 | .182 |
| *Group at 90°* | EB vs SC | 45 | −9.806 | 2.820 | −3.477 | [−15.33, −4.28] | < .001 | .002 |
| *Group at 90°* | LB vs SC | 46 | −5.625 | 2.848 | −1.975 | [−11.21, −0.04] | .048 | .096 |
| *MR within EB* | 90 vs 0° | 16 | −10.278 | 2.205 | −4.662 | [−14.60, −5.96] | < .001 | < .001 |
| *MR within EB* | 180 vs 0° | 16 | −13.425 | 2.205 | −6.089 | [−17.75, −9.10] | < .001 | < .001 |
| *MR within EB* | 180 vs 90° | 16 | −3.147 | 2.205 | −1.427 | [−7.47, 1.17] | .153 | .153 |
| *MR within LB* | 90 vs 0° | 17 | −7.895 | 2.139 | −3.691 | [−12.09, −3.70] | < .001 | < .001 |
| *MR within LB* | 180 vs 0° | 17 | −7.834 | 2.139 | −3.662 | [−12.03, −3.64] | < .001 | < .001 |
| *MR within LB* | 180 vs 90° | 17 | +0.061 | 2.139 | +0.029 | [−4.13, 4.25] | .977 | .977 |
| *MR within SC* | 90 vs 0° | 29 | −2.369 | 1.638 | −1.447 | [−5.58, 0.84] | .148 | .444 |
| *MR within SC* | 180 vs 0° | 29 | −1.696 | 1.638 | −1.036 | [−4.91, 1.51] | .300 | .601 |
| *MR within SC* | 180 vs 90° | 29 | +0.673 | 1.638 | +0.411 | [−2.54, 3.88] | .681 | .681 |

#### Text S7 – Correlation analyses

#### Table S14. Results from partial Pearson and Fisher r-to-z tests.

| *Test* | *Groups* | *n* | *r* | *z* | *p* | $p_{FDR}$ |
|---|---|---|---|---|---|---|
| | | *Partial Pearson Correlations* | | | | |
| *Pointing score × navigation performance* | EB | 16 | .746 | - | .002 | .002 ** |
| | LB | 17 | .832 | - | < .001 | < .001 *** |
| | SC | 29 | .775 | - | < .001 | < .001 *** |
| | All groups | 62 | .791 | - | < .001 | < .001 *** |
| *MR cost × navigation performance* | EB | 16 | .782 | - | < .001 | .003 ** |
| | LB | 17 | .528 | - | .043 | .065 † |
| | SC | 29 | .258 | - | .193 | .193 |
| | All groups | 62 | .495 | - | < .001 | < .001 *** |
| *Age × navigation performance* | EB | 16 | .175 | - | .533 | .533 |
| | LB | 17 | -.088 | - | .747 | .906 |
| | SC | 29 | -.435 | - | .021 | .041 * |
| *Age × pace* | EB | 16 | .233 | - | .404 | .533 |
| | LB | 17 | .180 | - | .506 | .906 |
| | SC | 29 | .379 | - | .047 | .062 † |
| *Age × pointing score* | EB | 16 | .410 | - | .129 | .516 |
| | LB | 17 | -.249 | - | .353 | .906 |
| | SC | 29 | -.522 | - | .004 | .018 * |
| *Age × MR cost* | EB | 16 | .235 | - | .399 | .533 |
| | LB | 17 | -.032 | - | .906 | .906 |
| | SC | 29 | -.096 | - | .628 | .628 |
| *BDI × navigation performance* | LB | 17 | .372 | - | .172 | .344 |
| *BDI × pace* | LB | 17 | -.159 | - | .572 | .572 |
| *BDI × pointing score* | LB | 17 | .479 | - | .071 | .282 |
| *BDI × MR cost* | LB | 17 | .215 | - | .441 | .572 |
| *SBSOD × navigation performance* | EB | 16 | .490 | - | .075 | .085 † |
| | LB | 13 | .609 | - | .047 | .154 |
| | All Blind (EB + LB) | 29 | .517 | - | .007 | .014 * |
| *SBSOD × pointing score* | EB | 16 | .771 | - | .001 | .005 ** |
| | LB | 13 | .554 | - | .077 | .154 |
| | All Blind (EB + LB) | 29 | .614 | - | .001 | .003 ** |
| *SBSOD × pace* | EB | 16 | -.477 | - | .085 | .085 † |
| | LB | 13 | -.124 | - | .717 | .717 |
| | All Blind (EB + LB) | 29 | -.379 | - | .056 | .056 † |
| *SBSOD × MR cost* | EB | 16 | .586 | - | .028 | .055 † |
| | LB | 13 | .215 | - | .526 | .701 |
| | All Blind (EB + LB) | 29 | .427 | - | .030 | .040 * |
| *Video games × navigation performance* | EB | 16 | .223 | - | .443 | .886 |
| | LB | 15 | .104 | - | .734 | .979 |
| | All Blind (EB + LB) | 31 | .163 | - | .406 | .786 |
| *Video games × pointing score* | EB | 16 | -.048 | - | .870 | .943 |
| | LB | 15 | .184 | - | .547 | .979 |
| | All Blind (EB + LB) | 31 | .052 | - | .794 | .794 |
| *Video games × pace* | EB | 16 | -.241 | - | .406 | .886 |
| | LB | 15 | .005 | - | .988 | .988 |
| | All Blind (EB + LB) | 31 | -.139 | - | .480 | .786 |
| *Video games × MR cost* | EB | 16 | .021 | - | .943 | .943 |
| | LB | 15 | -.319 | - | .288 | .979 |
| | All Blind (EB + LB) | 31 | -.106 | - | .590 | .786 |
| | | *Fisher r-to-z* | | | | |
| *Age × navigation performance* | EB vs LB | 33 | - | 0.66 | .509 | .678 |
| | EB vs SC | 45 | - | 1.83 | .067 | .134 |
| | LB vs SC | 46 | - | 1.11 | .268 | .684 |
| *Age × pace* | EB vs LB | 33 | - | 0.14 | .890 | .890 |
| | EB vs SC | 45 | - | -0.46 | .645 | .645 |
| | LB vs SC | 46 | - | -0.64 | .525 | .699 |
| *Age × pointing score* | EB vs LB | 33 | - | 1.72 | .085 | .340 |

| | | | | | | |
|---|---|---|---|---|---|---|
| | EB vs SC | 45 | - | 2.89 | .004 | .015 * |
| | LB vs SC | 46 | - | 0.95 | .342 | .684 |
| *Age × MR cost* | EB vs LB | 33 | - | 0.68 | .498 | .678 |
| | EB vs SC | 45 | - | 0.95 | .340 | .453 |
| | LB vs SC | 46 | - | 0.19 | .852 | .852 |

**Text S8 – Training procedures and exploration instructions**

Participants were familiarized to the tactile mazes and the virtual environment software by performing the with three mazes of increasing complexity: a T-maze, and two window mazes (see Fig. S7a-c). For this, they were guided by the experimenter to explore each maze, recognize their shape, recognize the different destinations.

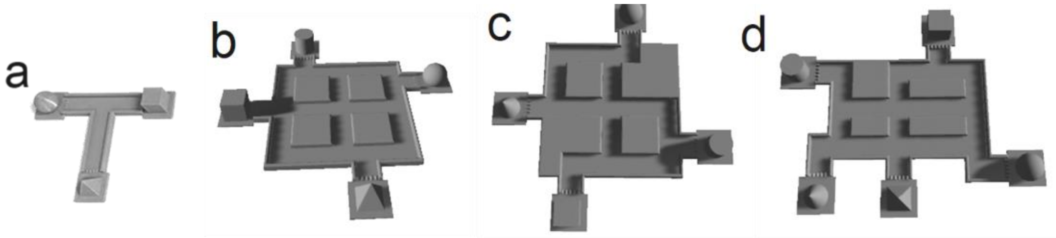


**Figure S7. Mazes used during the training session.** A. T-maze. B. Window-maze 1. C. Window- maze 2. (d) Final maze (main experiment, see main manuscript).

First, participants could tactilely explore and navigate freely in the T-maze version to get used to the sounds and the movement mechanics (see Text S1). Then, they proceeded to the task, with no time limits in either of the three phases. When participants were comfortable with the T-maze and were successful in at least two consecutive trials, they could proceed to the first window maze. For the window mazes, the experimenter repeated the same instructions for the exploration phase (see below) and proceeded to the task while still being able to exchange with the experimenter if they had questions or concerns.

**Exploration instructions (repeated for every maze, including the final maze in the main experiment):** [Before the exploration begins, the experimenter places the participant's dominant index on the central, southern-most destination] *Your index is now placed on the southern-most destination, don't move until the experiment begins and you hear "Maze. Beep,"* [see Movie S1]. *While exploring, try to learn the relative positions of each destination (for example, this one is more North, this one more East, compared to the other one, this one is more South-West). You also must learn where their doors are and which direction they are facing. This is important, because: in the pointing task, you'll have to imagine yourself like you are standing in the hallway, your back against the door (or as if you just stepped out that door and entered the hallway), you'll also start from that facing position at the beginning of the navigation trials. Furthermore, we want you to learn the layout of the maze, or its overall shape. How are the hallways connected? How and where do they cross? Are there main, long, vertical and/or horizontal hallways? How are the different destinations connected to these hallways. We don't want you to learn the specific routes between each destination, because as we will progress in the experiment, the number of destinations and hallways will grow, and learning routes will become increasingly difficult. The experiment will begin shortly* [the experimenter starts the virtual environment software, see Movie S1].

Participants experienced a simulated version of the task, but with extended time limits (unlimited time for tactile exploration: they told the experimenter when they were ready to proceed; 60s for navigation trials) and only three trials before the exploration phase returned. After the first tactile exploration period, they were asked to explain the layout of the maze and the experimenter could specify:

**Window maze 1 description:** *This maze has the shape of a window. The outer hallway, or perimeter, is a wide square of which all four sides have a destination attached to it. At the center of the maze, there are two crossing hallways (a + or cross shape) that can be shortcuts, for some cases. You can note that all shapes are not centered (or at the level of the cross), except one: the cube.*

When they could perform succesfully in at least three consecutive trials, they could proceed to the second window maze, with which they experienced a version of the task closer to the testing parameters (60 s exploration, 8 s ponting, and 30 s navigation trials). They were first explained:

**Window maze 2 description:** *This maze has the same window layout, but some corners are blocked. Therefore, you will be forced to go through the middle cross; this cross is now, in most cases, the main way to go from the complete North to complete South; or from the complete West to complete East; and vice versa* [The experimenter then repeats the exploration instructions].

Again, when they could succeed at three consecutive trials, they could proceed to the real version of the task with the final maze (Fig. S6d). with different mazes. Then, they could no longer interact with the experimenter.

**Text S9 – Psychosocial questionnaire**

**[Inclusion/Exclusion]**
Date of birth : ________________________________
Age at participation: _________________________
Sex (biologic): ________________________________
Gender identity : _____________________________
Blindness onset: __________________
Blindness cause : _____________________
Residual vision :

- Light perception? [ ]None, [ ]Low, [ ]strong
- Shape perception? [ ]None, [ ]Low, [ ]Strong
- Motion perception? [ ]None, [ ]Low, [ ]Strong

Condition progression (history): ___________________________
Other health conditions: ________________

**[Questionnaire, sociodemographic and other characteristics]**
Education (last level completed): _______________________________
Occupation: ________________________________________________
Dominant hand: _____________________________
Do you read Braille? [ ]Yes, [ ]No

- If yes, with which hand? [ ] left, [ ] right, [ ] both

At what age did you learn Braille? ___________
How long (in hours) do you read Braille in a day? _____________
How fast can you read Braille (words per minutes)? _____________
What is your primary mobility aid? [ ]Cane, [ ]Dog, [ ]Other:_______________
How many times do you travel outside your home in a day? In a week? In a month?
O&M lessons? [ ]Yes, [ ]No

- At what age did you have your first O&M lessons: ____________

Do you have experience with tactile maps? [ ]Yes, [ ]No

- If yes, how often do you use tactile maps? ___________________________

Do you have experience with videogames (or audio games)? [ ]Yes, [ ]No

- If yes, what type(s) of videogames, can you name examples? _____________

Santa Barbara Sense of Direction Scale (SBSOD) | Hegarty Spatial Thinking Lab | UC Santa Barbara
